# Multi-omics on truncating *ASXL1* mutations in Bohring Opitz syndrome identify dysregulation of canonical and non-canonical Wnt signaling

**DOI:** 10.1101/2022.12.15.520167

**Authors:** Isabella Lin, Angela Wei, Zain Awamleh, Meghna Singh, Aileen Ning, Analeyla Herrera, Bianca Russell, Rosanna Weksberg, Valerie A. Arboleda

## Abstract

*ASXL1* (*Additional sex combs-like 1*) plays key roles in epigenetic regulation of early developmental gene expression. *De novo* truncating mutations in *ASXL1* cause Bohring-Opitz syndrome (BOS, OMIM #605039), a rare neurodevelopmental condition characterized by severe intellectual disabilities, characteristic facial features, hypertrichosis, increased risk of Wilms tumor, and variable congenital anomalies including heart defects and severe skeletal defects giving rise to a typical ‘BOS posture’. These BOS-causing *ASXL1* variants are also high-prevalence somatic driver mutations in acute myeloid leukemia (AML). We use primary cells from BOS individuals (n = 18) and controls (n = 49) to dissect gene regulatory changes caused by *ASXL1* mutations using comprehensive multi-omics assays for chromatin accessibility (ATAC-seq), DNA methylation, histone methylation binding, and transcriptome in peripheral blood and skin fibroblasts. Our data shows that regardless of cell type, *ASXL1* mutations drive strong cross-tissue effects that disrupt multiple layers of the epigenome. The data showed a broad activation of canonical Wnt signaling at the transcriptional and protein levels and upregulation of *VANGL2*, a planar cell polarity pathway protein that acts through non-canonical Wnt signaling to direct tissue patterning and cell migration. This multi-omics approach identifies the core impact of *ASXL1* mutations and therapeutic targets for BOS and myeloid leukemias.

**Brief summary:** Germline *ASXL1* mutations that cause Bohring Optiz syndrome disrupt the epigenome and dysregulate gene expression resulting in activation of canonical and non-canonical Wnt signaling pathways.

## INTRODUCTION

*ASXL1* (Additional sex combs-like 1) is an essential protein in embryonic development (1). Eutherian mammals have three *Asx* homologs, *ASXL1, ASXL2,* and *ASXL3,* which share conserved domains; the ASX N-terminal (ASXN) domain with a HARE-HTH domain involved in putative DNA binding (2), the ASX homology (ASXH) domain with a characteristic LXXLL motif that mediates protein interactions and DEUBAD domain that activates BAP1, and the C-terminal plant homeodomain (PHD) important in “reading” post-translational histone modifications such as histone H3 Lys4 trimethylation (H3K4me3) (3–5) (**Figure 1A**). *ASXL1* plays a role in the modulation of the epigenetic landscape which regulates downstream transcription. The ASXH and PHD domains are highly conserved and are part of large epigenetic complexes, the Polycomb (PcG) complexes and the Deubiquitinase complexes. These epigenetic complexes direct genome-wide transcriptional regulation (3, 6, 7) through activation or repression of essential developmental genes. *Asx,* the *Drosophila melanogaster* ortholog, plays a critical role in anterior-posterior patterning (8) and mutant embryos exhibit incomplete head involution and posterior directed transformations of all abdominal segments (8). In mammals, *ASXL1* is essential in early neural crest (9), cardiac (10) and hematopoietic development (11, 12).

**Figure 1.**
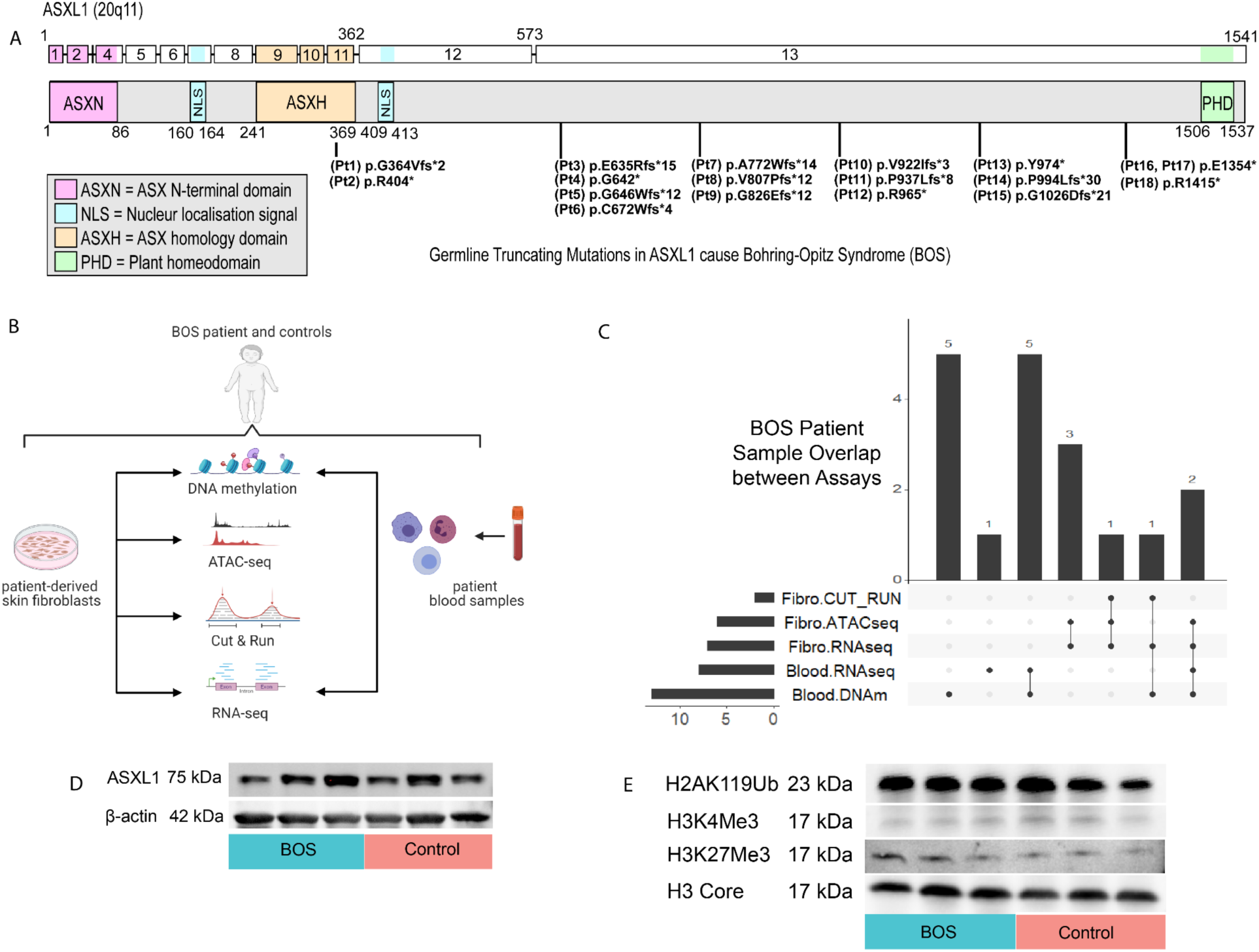
Multiomics study design for Bohring-Opitz Syndrome (BOS) caused by pathogenic mutations in *ASXL1*. **(A)** Schematic representation of the ASXL1 transcript (ENST00000375687.10) and protein (GenBank: ASXL1; NM_015338.6; GRCh37), its functional domains, and mutations causing BOS. Mutations listed correspond to patients in this study and are tagged with a Pt identifier. **(B)** Peripheral blood and dermal fibroblasts were collected from BOS patients and controls. Dermal fibroblast samples underwent epigenomic assays for assay for transposase-accessible chromatin with high-throughput sequencing (ATAC-seq), cleavage under targets & release using nuclease (CUT&RUN) and global transcriptome analysis using RNA sequencing (RNA-seq). Whole blood samples underwent DNA methylation analysis and global transcriptome analysis by RNA-seq. **(C)** Across the multi-omics assays and two specimen types we had varying degrees of overlap among assays. Six patients had one blood assay, five patients had both blood assays, four patients had multiple fibroblast assays, and three patients had data from assays across both specimen types. This totalled eighteen patients. **(D)** Western blot for representative BOS (n=3) and representative control (n=3) fibroblast whole cell lysate extracts shows no significant difference in total ASXL1 protein from ImageJ quantification. **(E)** Western blot for representative BOS (n=3) and representative control (n=3) fibroblast histone extracts shows no significant difference in H2AK119 ubiquitination, trimethylation of H3K4 and trimethylation of H3K27me3 from ImageJ quantification.

ASXL1 directs histone modifications through the Polycomb Repressive Complex 1 and 2 (PRC1/2) and the Polycomb Repressive Deubiquitinase complex (PR-DUB). PRC1 and 2 control activation of developmental stemness and differentiation, including progressive restriction of neural progenitor cell multipotency and production of mature cortical neurons during corticogenesis (13). This control of the stem-cell state occurs through monoubiquitination at histone H2A lysine 119 (H2AK119ub) via PRC1 and methylation at histone H3K27 via PRC2 (3, 14, 15). Together, these complexes regulate key signaling pathways such as Wnt signaling in tissue-specific and developmental contexts, with PRC2 accessory proteins regulating Wnt signaling during erythropoiesis (16) and PRC1 regulating the Wnt/β-catenin pathway through a positive feedback loop in hepatocellular carcinoma (HCC) (17).

The PR-DUB, which includes ASXL1 that binds BRCA1-associated protein 1 (BAP1) (18–20), plays key roles in brain development (21–23) and regulation of myeloid differentiation through H2AK119 deubiquitination (19, 24). While it is clear that *ASXL1* protein truncating mutations disrupt core developmental processes across multiple organ systems, its role as an essential chromatin modifier in human development has not been fully elucidated.

### *ASXL1* mutations in Bohring Opitz Syndrome (BOS) and in hematologic malignancies

*De novo* protein truncating mutations of *ASXL1* cause a rare genetic disorder, Bohring-Opitz Syndrome (BOS, OMIM #605039). BOS is characterized by profound intellectual disability, developmental delay, seizures, variable anomalies including heart defects, higher risk of Wilms tumor, and ‘BOS posture’ (25, 26). As of 2018, only 46 clinically diagnosed individuals have been reported in the literature, with less than half (20/46) molecularly confirmed (25). Pathogenic truncating mutations in *ASXL1* are enriched in exons 12 and 13, the penultimate and ultimate exons (3), resulting in premature truncation of the highly conserved C-terminal PHD domain. Intriguingly, truncation mutations in ASXL1 homologs *ASXL2* (Shashi-Pena Syndrome, OMIM#6171901) (27) and *ASXL3* (Bainbridge-Ropers Syndrome, OMIM #615485) (23, 28) are not enriched or clustered in the homologous C-terminal region despite sharing multiple conserved domains (3).

The same protein-truncating mutations causing BOS are also observed as somatic driver mutations in myeloid malignancies such as chronic myelomonocytic leukemia (CMML, ∼45%), myelodysplastic syndromes (MDS, 16%), myeloproliferative neoplasms (∼10%) and acute myeloid leukemia (AML, secondary 30%, *de novo* 6.5%) (29–32). Studies have identified low-frequency somatic mutations in *ASXL1* correlated with mutagenic processes and increasing age (33), defining a new pre-malignancy state called clonal hematopoiesis of indeterminate potential (33, 34). *ASXL1* regulates the delicate interplay between proliferation and differentiation of stem progenitor cell populations (9, 35) and therefore, both germline and somatic *ASXL1* mutations disrupt the proliferation-differentiation balance and promote stem-cell identity over differentiation in BOS and myeloid leukemias.

Recent *ASXL1* functional studies have been carried out as transgenic overexpression of mutant and wild-type ASXL1 proteins in cell-line systems. These studies are difficult to interpret because they are not reflective of endogenous levels or cell-type specific functions of ASXL1 or they are in non-human model systems (36). Given the multiple essential functions of ASXL1 as part of PRC1 and PRC2 in neural progenitor multipotency (13), and key role of PR-DUB in brain development (21–23), we hypothesize that protein truncating mutations of *ASXL1* dysregulate global transcription, cellular homeostasis and downstream signaling pathways.

The broad phenotypic effects of *ASXL1* mutations in BOS patients and myeloid malignancies suggest that *ASXL1* drives essential and core gene regulatory features across early development and disease. We hypothesized that *ASXL1* mutations share common cellular effects that might therefore be detectable as broad and high-effect epigenomic and transcriptomic signatures across cells and tissues. We also examined whether BOS cells harbored decreased mRNA or protein levels since protein truncating *ASXL1* mutations occur in the last two exons of the gene, exons 12 and 13, and are predicted to escape nonsense mediated decay (37).

To study BOS molecular pathogenesis, we collected samples from 18 BOS patients, one of the largest ASXL rare disease cohorts published, and performed multi-omics analysis. Our approach circumvents the confounding variable of cancer-derived cell models that harbor multiple additional karyotypic and genomic mutations (31, 38) and focuses our study on the singular effect of pathogenic *ASXL1* mutations. This integrated multi-omics study characterizes the impact of *ASXL1* truncating mutations on the epigenome and transcriptome and our work shows for the first time that *ASXL1* mutations aberrantly activate Wnt signaling pathway and non-canonical Wnt planar cell polarity (PCP) genes.

The canonical Wnt signaling is an evolutionarily-conserved signaling pathway that is essential for development (39), and plays important roles in stem cell biology and regulation of hematopoiesis (40). In the Wnt-active state, Wnt ligands bind to the transmembrane receptor Frizzled (FZD) and stimulates the co-receptor Low Density Receptor-Related Protein (LRP)5/6, which play critical roles in initiation of Wnt signaling transduction (41). Activation of FZD and LRP5/6 recruits and inactivates the “destruction complex” that targets phosphorylation, ubiquitination and degradation of β-catenin through the proteasome. Inactivation of the “destruction complex” occurs through dissociation of glycogen synthase kinase 3 beta (GSK3β) from AXIN, and concurrent inactivation of the default quiescent-state “β-catenin destruction complex” (42–44). This means that β-catenin cannot be phosphorylated, and thus cannot be degraded (45, 46). The activation of the canonical Wnt pathway results in cytoplasmic accumulation and subsequent nuclear translocation of β-catenin which, along with T-cell factor/lymphoid enhancer factor (TCF/LEF) transcription factors transcriptionally co-activates Wnt-target genes (47).

To examine cross-omics dysregulation in more depth, we identified *Van gogh-like 2* (*VANGL2)*, a non-canonical Wnt and planar cell polarity (PCP) protein, which was highly dysregulated across all the -omics assays conducted across blood and fibroblast BOS samples. *VANGL2* regulates polarized cellular migration and differentiation and tissue morphology during development. In the nervous system, the PCP pathway regulates neuronal maturation and has a functional role in neural complex formation (48). VANGL2 intersects with the canonical pathway through activation of DVL which activates non-canonical pathways and drives c-JUN mediated expression. We hypothesize that upregulation of *VANGL2* disrupts canonical and non-canonical signaling pathways and underlies the clinical phenotypes observed in patients with BOS and other mutations in ASXL. To date, no studies have established a link between *ASXL1* disorders and dysregulation of Wnt-signaling.

In this study, we identify and test the mechanisms by which truncating *ASXL1* mutations in BOS may dysregulate the canonical and non-canonical Wnt signaling pathways. These findings elucidate novel regulatory mechanisms underlying BOS pathogenesis.

## RESULTS

### Study Design

We recruited 18 individuals with BOS (**Supplemental Table 1**), 23 sex-matched and genetically related controls, and 26 sex- and age-matched controls (**Supplemental Table 2**). This is the largest cohort of BOS patients studied at the molecular level. All BOS individuals have clinically identified *ASXL1* mutations in the last two exons of the gene that are predicted to cause protein truncating variants (NM_015338.6) between amino acids 364 to 1415 (**Figure 1A**, **Supplemental Table 1**). All mutations were verified through bulk RNA-seq data (**Supplemental Figure 1**) to confirm specimen identity and a summary of patient demographics are detailed in **Supplemental Table 3**. Clinical data of this cohort are available through the ASXL registry, and a subset of patients were previously reported by Awamleh et al. (49).

Consented individuals donated peripheral blood, a skin punch biopsy or both. DNA and RNA was extracted from whole blood (n = 14) and peripheral blood mononuclear cells (PBMCs) were cryopreserved. Skin punch biopsies (n = 8) were processed to obtain dermal fibroblasts. We obtained both sample types from 4 of the 18 BOS individuals in this study which allowed us to integrate data across sample types to identify tissue-dependent and -independent dysregulation caused by truncating *ASXL1* mutations. We next conducted a comprehensive multi-omics analysis across BOS patient-derived peripheral blood and dermal fibroblasts **(Figure 1B)**. We assessed global changes to the epigenome using three approaches: chromatin accessibility using assay for transposase-accessible chromatin using sequencing (ATAC-seq) (50), H3K4 trimethylation and H3K27 trimethylation enrichment using cleavage under targets and release using nuclease (CUT&RUN) (51), and the DNA methylation landscape through Illumina Epic Arrays (49). Not all samples were assessed by all methods and therefore the overlap between assays, blood samples (n = 14) and fibroblast samples (n = 8) from BOS individuals is outlined in **Figure 1C.** We derived disease-specific DNA methylation (DNAm) signatures (49) and transcriptional signatures from RNA-seq using blood and fibroblast cells (**Figure 1B)**.

*ASXL1* is expressed across multiple cell types (**Supplementary Table 4**), which is consistent with the fact that the clinical phenotype spans multiple organ systems. To test our hypothesis that *ASXL1* mutations may have high-level epigenomic and transcriptomic signatures across affected cells and tissues, we conducted RT-qPCR using *ASXL1* primers (**Supplemental Table 5**). We found no significant differences in *ASXL1* expression levels between BOS and control samples (**Supplemental Figures 2A and 2B**).

We next asked if the truncating mutation caused differences in wild-type ASXL1 protein levels due to decreased translational efficiency or stability of the truncated allele. We identified a high quality antibody by testing multiple, commercially-available antibodies (**Supplemental Table 6**, **Supplemental Figure 3A**) in CACO2 cells, a cell line with very high *ASXL1* RNA expression (**Supplemental Table 4**), and HEK293T cells transfected with FLAG-tagged truncated *ASXL1* (52). We identified two antibodies (ab228009, 12F9) that showed bands at the same molecular weight of 75 kDA, one of which (ab228009) showed higher expression of ASXL1 in CACO2 cells and overlapped with ASXL1-construct with high-affinity FLAG antibody (**Supplemental Figure 3B**). Western blot for ASXL1 identified no differences in total ASXL1 protein levels between patient and control fibroblasts (**Figure 1D**).

Finally, we extracted histones from fibroblast cells and quantified global changes for histone modification. The mutations in BOS terminate the protein before the PHD domain, a histone- or DNA-binding domain reported to recognize a subset of histone modifications including H3K4me and H3K27me3 (5), and are thought to affect levels of H3K27me3 and H2AK119ub (19). We performed histone immunoblotting for these specific modifications and do not observe any global changes to H3K4me3, H3K27me3, or H2AK119Ub (**Figure 1E**, **Supplemental Figure 4**)

### Transcriptomic analysis identifies transcriptional pathways disrupted in BOS

One of the core functions of *ASXL1* is to activate the epigenome to express certain genes and pathways. Changes to the epigenome activate or repress transcription and RNA-seq can identify the effects of the transcriptional rewiring due to epigenetic mutations. To address this, we performed RNA-seq in blood and dermal fibroblasts from individuals with BOS. Transcriptomic data was generated for a total of 33 RNA-seq libraries: 8 BOS-blood, 7 BOS fibroblasts, 11 control blood, and 7 control fibroblasts (**Supplemental Table 7 and 8**, **Methods**). Similar to the qRT-PCR data, *ASXL1* normalized gene expression was not significantly different between BOS and control samples (**Supplemental Figures 2C and 2D**). We then examined whether the *ASXL1* reference and mutant alleles were equally represented in blood (**Figure 2A**) and fibroblast (**Figure 2B**) RNA-seq data. This was performed by calculating the variant allele frequency (VAF) at the mutation site across all the BOS samples (**Supplemental Figure 5**). For germline disorders, we would expect the VAF of the mutant allele to represent between 30 to 70 percent of the reads covering the mutation site (0.3 < VAF < 0.7). Most BOS samples fell within the expected range (**Supplemental Figures 2A and 2B**, **Supplemental Table 9**), suggesting that these late-truncating alleles escape nonsense-mediated decay. One key exception was Pt6, with truncating mutation at amino acid 672 who had undergone treatment with actinomycin D and vincristine for Wilms tumor years prior to sample collection. In fibroblast RNA-seq, this sample had an *ASXL1* allele ratio of 36.9% pathogenic allele, which is within expected germline levels. However, in the same individual’s blood RNA-seq, 100% of reads (27/27) over the *ASXL1* mutation contained the pathogenic mutation and no reference allele was identified. This loss of heterozygosity may have occurred as a selective growth advantage during tumorigenesis, during treatment or could represent new clonal hematopoiesis.

**Figure 2:**
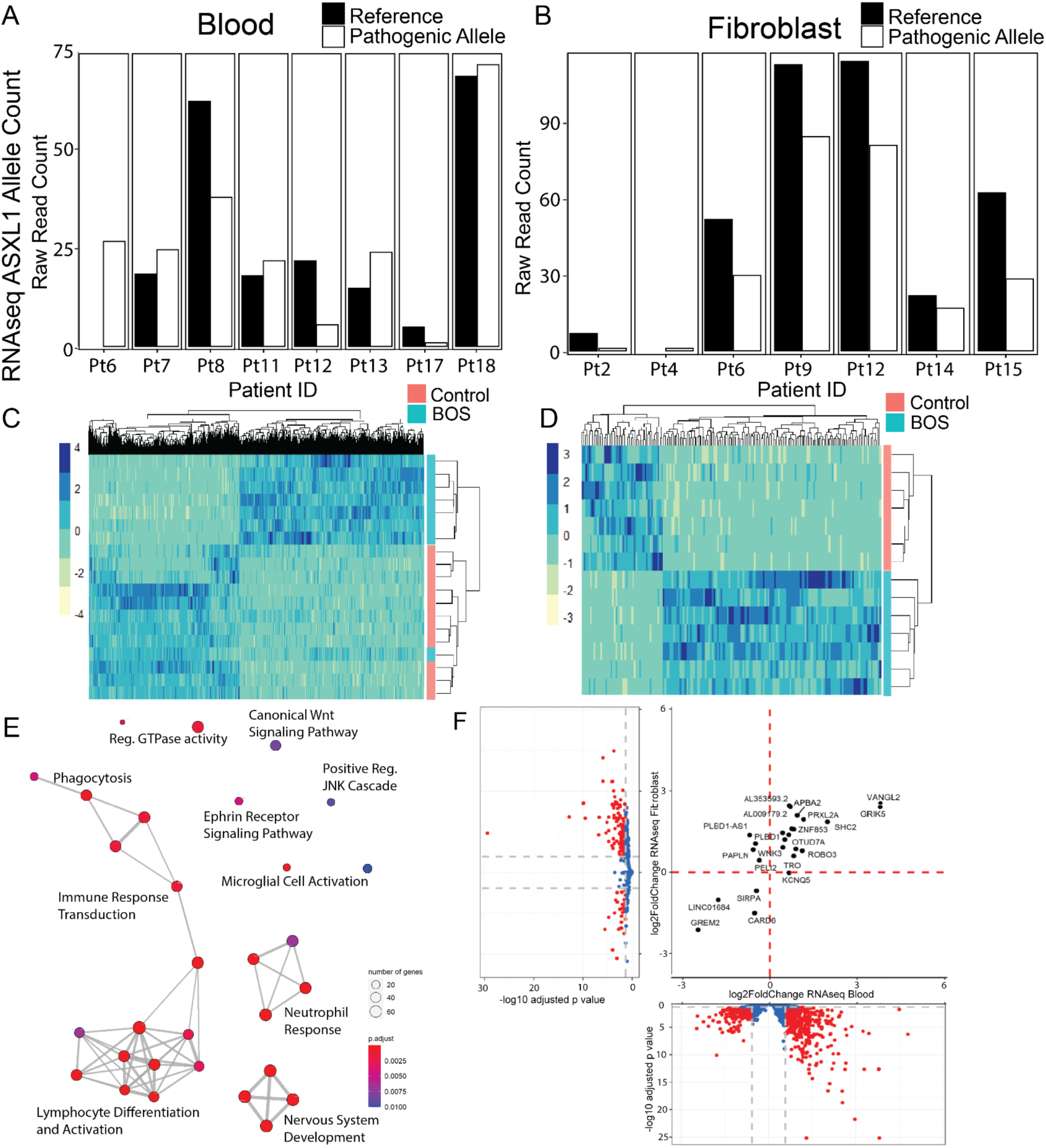
Pathogenic mutations in *ASXL1* cause tissue-specific and tissue-independent effects on gene expression. **(A)** RNA sequencing (RNA-seq) shows raw read counts for the *ASXL1* reference allele (black) and pathogenic allele (white, with black outline) at each BOS patient’s respective mutation in blood and **(B)** patient-derived fibroblast samples. **(C)** RNA-seq heatmap of all significant differentially expressed genes (DEGs) with adjusted p < 0.05 and |log_2_FC| ≥ 0.58 between BOS (blue) and controls (pink) show 1097 DEGs in blood and **(D)** 155 DEGs in fibroblasts, with 590/1097 (53.8%) and 125/155 DEGs (80.6%) being more upregulated in BOS patients respectively. **(E)** Gene ontology of all significant DEGs in blood RNA-seq highlights enrichment of genes related to nervous system development and canonical Wnt signaling pathway. **(F)** Volcano plots for BOS compared to control RNA-seq in blood (x-axis) and fibroblast (y-axis) identifies a core subset of 25 shared dysregulated transcripts, with 21/25 DEGs (84%) dysregulated in the same direction.

To assess the pathogenesis of truncating *ASXL1* mutations, we wanted to examine the effects on global transcription across different tissues. *ASXL1* is ubiquitously expressed across the body at low levels, and is more highly expressed in skin (32.55 TPM) than in whole blood (8.03 TPM) (**Supplemental Table 4**). We expected a much higher number of significant differentially expressed genes (DEGs) in blood RNA-seq than in fibroblast RNA-seq because blood RNA represents a bulk assessment of heterogeneous cell types which can introduce an extra source of differential expression. We see significantly more gene expression dysregulation in blood with 2118 significant DEGs (**Figure 2C**) compared to fibroblast RNA-seq with 177 significant DEGs (**Figure 2D**). Complete lists of DEGs in each dataset are in **Supplemental Table 10** for BOS fibroblast and **Supplemental Table 11** for BOS blood. After filtering for fold change (|log_2_FoldChange| ≥ 0.58), we see 1097 significant DEGs in blood and 155 significant DEGs in fibroblasts.

Both RNA-seq analysis of blood and fibroblasts identified larger proportions of DEGs that are upregulated in BOS patients than in controls. In blood, 590/1097 DEGs (53.8%) were upregulated (**Figure 2C**), and in fibroblast cells 125/155 DEGs (80.6%) (**Figure 2D**). In blood, top DEGs included *GRIK5* (*Glutamate receptor, Kainate 5,* log_2_FC 3.8), *VANGL2* (*Vang-like protein 2,* log_2_FC 3.8), and *GREM2* (*Gremlin 2*, BMP antagonist, log_2_FC -2.5). Top DEGs in the fibroblast RNA-seq dataset include *UGT3A2* (log_2_FC 4.8), *VANGL2* (log_2_FC 2.5), *GRIK5* (log_2_FC 2.5) and *GREM2* (log_2_FC -2.1). Gene ontology analyses were performed using the list of significant DEGs. Despite neither tissue being neural in origin, *regulation of neuron projection development* (p_adj_ = 9.82 x 10^-5^ in blood, p_adj_ = 0.02 in fibroblasts) was identified as a gene ontology biological process enriched in both BOS tissues (**Figure 2E, Supplemental Tables 12 and 13**). These enrichments were driven by genes that play key roles in early morphogenesis, particularly neurodevelopmental processes.

Our findings from gene ontology analyses in blood and fibroblast also identified some gene expression changes that are tissue-specific. Blood DEGs were enriched for hematological processes such as T-cell activation (p_adj_ = 3.23 x 10^-8^), neutrophil activation (p_adj_ = 1.90 x 10^-5^), axogenesis (p_adj_ = 1.90 x 10^-5^), leukocyte cell-cell adhesion (p_adj_ = 3.62 x 10^-5^) (**Figure 2E, Supplemental Table 12**). Fibroblast DEGs were enriched for structural cell processes such as dysregulation of potassium ion transport (p_adj_ = 0.004) and regulation of membrane potential (p_adj_ = 0.02) (**Supplemental Table 13**).

To examine potential common, cross-tissue effects of truncating *ASXL1* mutations, we correlated all our BOS blood RNA-seq DEGs and all our BOS fibroblast RNA-seq DEGs to identify shared dysregulated transcripts (**Figure 2F**). We identified a core subset of 25 genes dysregulated across both tissue types in the same direction, suggesting a strong tissue-independent *ASXL1* effect that supersedes tissue type. Notable genes include *VANGL2, GRIK5* and *GREM2*.

Motif enrichment analysis of our RNA-seq fibroblast data was performed using HOMER (Hypergeometric Optimization of Motif EnRichment)(53). Both known motifs and *de novo* methods identified significant enrichment of the following transcription factor binding sites: RARa (p_adj_ = 1 x 10^-2^, BOS 34.38%, background 27.61%), FOXO3 (p_adj_ = 1 x 10^-2^, BOS 8.44%, background 5.04%), FOXK2 (p_adj_ = 1 x 10^-2^, BOS 8.12%, background 4.88%) and ERRg (p_adj_ = 1 x 10^-2^, BOS 13.12%, background 9.03%). Pathway analysis for the enriched motifs in BOS fibroblasts identified roles of these motifs in leukemia/lymphoma pathways (p_adj_ = 5.9 x 10^-8^) and neurogenesis (p_adj_ = 1.7 x 10^-7^) (**Supplemental Table 14**) which is surprising given the cell type.

### Differential chromatin accessibility in BOS allows for aberrant activation of developmental and morphogenic pathways

To correlate gene expression profiles with chromatin accessibility (**Figure 3A**), ATAC-seq was performed on BOS and control fibroblast cells. Chromatin accessibility, identified by increased reads over a genomic region in ATAC-seq, is positively correlated with gene expression (50, 54).

**Figure 3:**
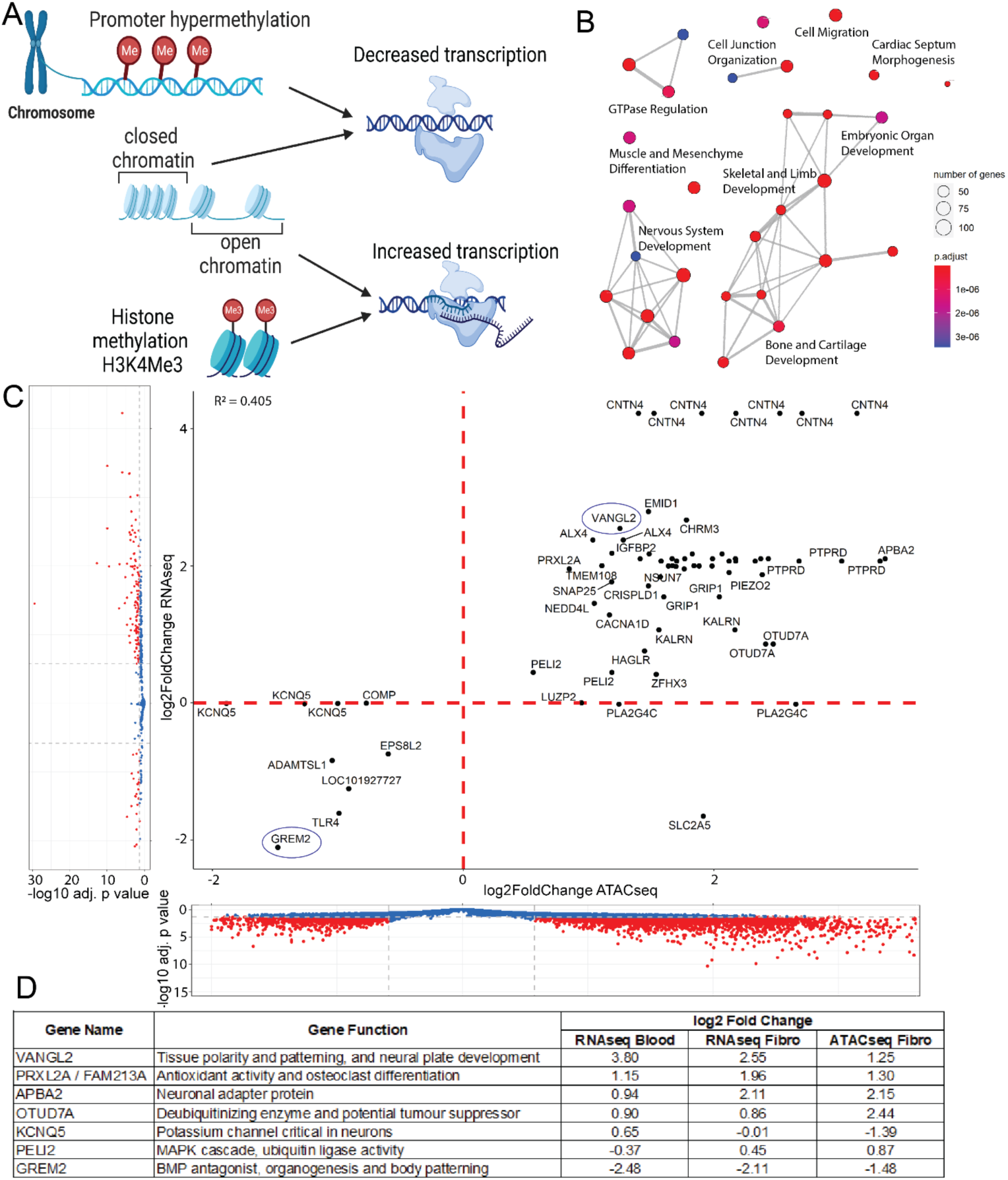
Epigenetic dysregulation in BOS patient-derived fibroblasts drive transcriptomic dysregulation. **(A)** Promoter hypermethylation and closed chromatin are associated with decreased transcription while activating histone methylation such as H3K4Me3 at promoters and open chromatin increase transcription. **(B)** Gene set enrichment of all significant (Bonferroni adjusted p<0.05) DEGs using ATAC-seq identified key dysregulated pathways. **(C)** Integration of chromatin accessibility (ATAC-seq, x-axis) and gene expression (RNA-seq, y-axis) in BOS patient fibroblasts identified a strong positive correlation (R^2^ = 0.405) between chromatin accessibility and gene expression. We identified a set of 37 common dysregulated transcripts across the epigenome and transcriptome (right). DEGs were considered significant (red) for |log_2_FC|>0.58 and adjusted p-value > 0.05. **(D)** Integration of datasets across tissue types and across sample types identifies a subset of overlapping genes that are significantly dysregulated in BOS. These 7 genes were identified across blood and fibroblast RNA-seq as well as fibroblast ATAC-seq. Gene functions and their respective significant log_2_FC are shown.

All ATAC-seq libraries had the expected distribution of fragment lengths, with an average of 52% of fragments being small (<200 bp length) representing open chromatin regions, and progressively fewer fragments of larger size spanning nucleosomes (>300 bp length) (**Supplemental Figure 6**, **Supplemental Table 15**). Of note, BOS patient fibroblasts had a mean insert size of 184bp, representing more open chromatin regions, while control fibroblasts had a median insert size of 242bp, representing less open chromatin regions. PCA identified clear separation of BOS and control samples on PC1 (47% variance) (**Supplemental Figure 7A**). 4246 significant peaks (p_adj_ > 0.05, abs(log_2_FC) > 0.58) corresponding to differentially accessible regions were identified, with 3036 peaks (70.02%) more differentially open and 1300 peaks (29.98%) more closed in BOS patients compared to controls (**Supplemental Figure 7B**, **Supplemental Table 16**). This mapped to a total of 3054 unique genes. Top differentially accessible peaks mapped to *FBXL20* (log_2_FC -8.3), *LINGO1* (log_2_FC -7.2), *RUNX3* (log_2_FC 4.4) and *CTNNB1* (log_2_FC 3.8).

We identified gene set enrichments in multiple key developmental systems including muscle development and differentiation (p_adj_ = 9.11 x 10^-13^), skeletal system development (p_adj_ = 1.0 x 10^-9^), regulation of neurogenesis (p_adj_ = 1.35 x 10^-9^), limb morphogenesis (p_adj_ = 1.38 x 10^-8^), renal system development (p_adj_ = 2.14 x 10^-8^), and cardiac septum morphogenesis (p_adj_ = 2.67 x 10^-8^) (**Figure 3B**, **Supplemental Table 17**). Key motifs that were identified in the differentially accessible chromatin regions of our ATAC-seq data suggest core factors that drive dysregulation of the DEGs. Motif enrichment analysis of our ATAC-seq data using both known motif and *de novo* methods identified significant enrichment of the following transcription factor binding sites: JunB (p_adj_ = 1 x 10^-208^, BOS 18.43%, background 5.23%), RUNX1 (p_adj_ = 1 x 10^-203^, BOS 26.45%, background 10.09%) and Fra1 (p_adj_ = 1 x 10^-196^, BOS 17.44%, background 4.95%) (**Supplemental Table 18**).

### Chromatin accessibility drives changes in gene expression in BOS

To determine whether some of the transcriptional dysregulation identified in BOS occurs through chromatin accessibility, we integrated DEGs from fibroblast RNA-seq with differentially accessible chromatin regions identified through ATAC-seq. This revealed a strong positive correlation (R^2^ = 0.405) between increased chromatin accessibility and increased gene expression of the same gene (**Figure 3C**). 71 differentially accessible chromatin regions aligned to 37/117 unique DEGs (20.9%) in the fibroblast RNA-seq dataset (**Supplemental Table 19**). A subset of these peaks mapped to the promoter-TSS (transcriptional start site) region, which usually indicate stronger effects on transcription. 14/71 (19.7%) peaks mapped to promoter-TSS regions, including the differentially accessible peak for *VANGL2,* corresponding to 13/37 (35.1%) unique DEGs in the fibroblast RNA-seq data (**Supplemental Figure 8**). These peaks showed a clear association of increased chromatin accessibility and increased transcription.

A subset of the DEGs identified through ATAC-seq translated across tissue types and were also found to be dysregulated in RNA-seq of BOS fibroblast and blood samples (**Figure 3D**). We identified seven genes that were significant and differentially expressed across omics levels (ATAC-seq and RNA-seq) and across tissues (blood and fibroblasts); *VANGL2, PRXL2A, APBA2, OTUD7A, KCNQ5, PELI2* and *GREM2* (a representative selection shown here in **Supplemental Figures 9 - 11**). These genes play significant roles in body patterning, neuron function, neural plate development and ubiquitination among other functions.

### Truncating *ASXL1* mutations dysregulate DNA methylation, and contribute to changes in gene expression

Endogenous CpG methylation levels at promoter regions are negatively correlated with gene expression (54) (**Figure 3A**). Thus, increased DNA CpG methylation or more closed chromatin are correlated with lower gene expression. To generate a BOS-specific DNA methylation (DNAm) signature, we profiled genome-wide DNAm in blood from BOS individuals (n=14) compared to 26 sex- and age-matched control subjects(55). DNA methylation identified 8596 differentially methylated CpG sites (FDR < 0.05) associated with an ENSEMBL gene, with 5773 CpG sites overlapping genes and mapping to 3803 unique genes. 763 differentially methylated CpG sites met a threshold of |Δβ| > 0.10 (10% DNAm difference) using linear regression modeling (**Supplemental Table 20**).

To examine the likelihood of statistically significant differential methylation resulting in downstream transcriptional effects, we correlated BOS blood DNA methylation and RNA-seq expression profiles. Endogenous CpG methylation levels at promoter regions are negatively correlated with gene expression (54) (**Figure 3A**). We first identified differentially methylated CpG sites (FDR < 0.05) and DEGs from RNA-seq (padj < 0.05) separately. We found 672 differential CpG sites that mapped to 341 unique DEGs in the blood RNA-seq dataset, with 143 genes represented by more than one differential CpG site (**Supplemental Table 21**). We filtered for highly differentially methylated sites (|Δβ| > 5%) and highly differentially expressed transcripts (|log_2_FoldChange (log_2_FC)| > 1.5). After filtering, we retained 50/672 CpG sites (7.44%) which corresponded to 24/341 unique genes (7.04%) (**Figure 4A**). 11 genes were represented by >1 CpG site, including *PSMA8* with 8 CpG sites and *GRIK5* with 2 CpG sites. *VANGL2* was represented by 1 CpG site. The correlation between these DNA methylation differences and transcriptomic changes indicates a steady-state effect of dysregulated DNA methylation on gene expression.

**Figure 4:**
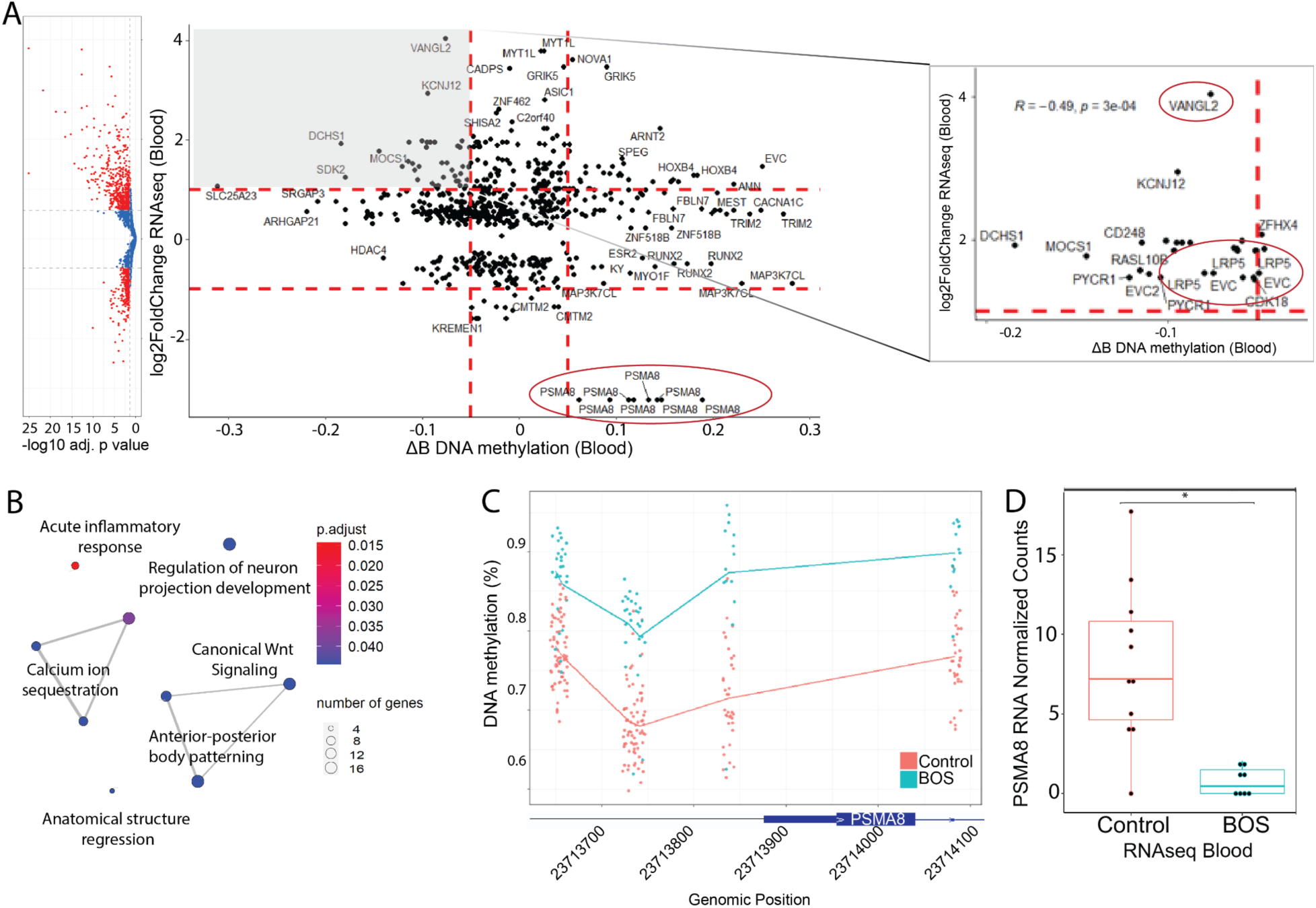
DNA methylation drives transcriptomic dysregulation in BOS blood and fibroblasts identify common dysregulated transcripts enriched in Wnt signaling genes. **(A)** Integration of BOS patient blood samples across DNA methylation and RNA transcriptomic dysregulation identifies 672 differentially methylated CpG sites (adjusted p < 0.05) that correlate to 341 dysregulated transcripts in RNA-seq (adjusted p < 0.05). These significant DEGs were further filtered for RNA-seq |log_2_FC| ≥ 0.58, and DNAm |Δβ| ≥ 0.05, shown by the dotted red lines. After filtering, we retained 50/672 CpG sites (7.44%) and 24/341 unique genes (7.04%). Inset window shows common DEGs that are hypomethylated and transcriptionally upregulated, including *Vangl2*, and *LRP5*. **(B)** Analysis of enriched biological processes using this integrated set of dysregulated genes identified canonical Wnt signaling, anterior-posterior body patterning, regulation of neuron projection development and other biologically relevant pathways. Enrichment of Wnt signaling is exemplified by the dysregulation of downstream target *PSMA8*. **(C)** In BOS patients, *PSMA8* is hypermethylated in blood DNA methylation across 8 CpG sites (Δβ 6.1% to 18.9%) and **(D)** shows strong downregulation in blood RNA-seq (log_2_FC -2.92).

### Gene set enrichment analysis of DNA methylation and RNA sequencing identifies activation of canonical Wnt pathway signaling in BOS samples

The overlapping findings from the DNAm and RNA-seq datasets prompted us to run gene ontology analysis for common gene targets, using two complementary methods, GREAT (56) and clusterProfiler *v3.12.0* (*57*). We filtered for gene ontologies with p_adj_ < 0.05 for significance. In both analyses, gene set enrichments were identified in the Wnt signaling pathway (GO#0060070, 17 genes, p_adj_ = 0.045), anterior-posterior pattern specification process (GO#0007389, 19 genes, p_adj_ = 0.045), and regulation of neuron projection development (GO#0010975, 19 genes, p_adj_ = 0.045), among other biologically relevant pathways (**Figure 4B**, **Supplemental Table 22**). Key genes that are enriched for these pathways include *PSMA8, WNT7A, FZD3, LRP5* and *LRP6* in canonical Wnt signaling and body pattern specification. The latter pathway enrichment was also driven by strong dysregulation in *VANGL2.* These overlapping gene targets and pathways between methylome and transcriptome in BOS patients suggest strong, coordinated, epigenetically-driven effects of truncating *ASXL1* mutations.

Three differentially methylated CpG sites, within the TSS, were identified for *Ellis van Creveld* (*EVC*) and *EVC2*. *EVC* and *EVC2* were both hypomethylated at the TSS (-5% and -3.5% respectively) and transcriptionally upregulated (log_2_FC 1.50 and 1.54 respectively) in BOS patient blood.

*Proteasome 20S subunit alpha 8* (*PSMA8*) is a key driver gene in the canonical Wnt-signaling enrichment identified in blood DNA methylation and RNA-seq data. We identified significant hypermethylation of *PSMA8* across all 8 CpG sites identified across the promoter and TSS region with Δβ from 6.1% to 18.9% (**Figure 4C, Supplemental Figure 12A - 12F**). *PSMA8* was also downregulated at the transcriptional level in BOS patient blood (log_2_FC -2.92, **Figure 4D**). *PSMA8* was not expressed in BOS or control fibroblasts, with normalized read counts of 0 in all samples, and a read count of 1 in one BOS fibroblast sample (**Supplemental Figure 12G**).

### Truncating *ASXL1* mutations disrupt canonical and non-canonical Wnt signaling

Through integration of these data, we identified dysregulation of the canonical Wnt pathway in gene set enrichment analysis of RNA-seq (**Figure 2E**) and DNA methylation studies (**Figure 4B**), driven by a consistent pattern of canonical Wnt/β-catenin pathway upregulation in the RNA-seq DEGs (**Supplemental Table 23**). Briefly, we identified upregulation of Wnt ligands: *WNT1* (log_2_FC 0.87), *WNT7A* (log_2_FC 0.97), and *WNT10B* (log_2_FC 0.53) and upregulation of Wnt ligand receptor, *Frizzled* receptor *FZD3* (log_2_FC 0.87).

DNA methylation studies identified aberrant hypomethylation of *LRP5* (**Figure 5B**) and *LRP6* (**Figure 5E**), which are Wnt signaling co-receptors. We identified 4 hypomethylated DNAm sites for *LRP5* (Δβ -3.5% to -8.0%), 2 of which reside in an important regulatory CpG island, and 2 hypomethylated DNAm sites for *LRP6* (Δβ -2.7% to -4.0%), both of which reside in an important regulatory CpG island. We also identified related transcriptional upregulation of *LRP5* (log_2_FC 1.64, **Figure 5C and 5D**) and *LRP6* (log_2_FC 1.63, **Figure 5F and 5G**).

**Figure 5:**
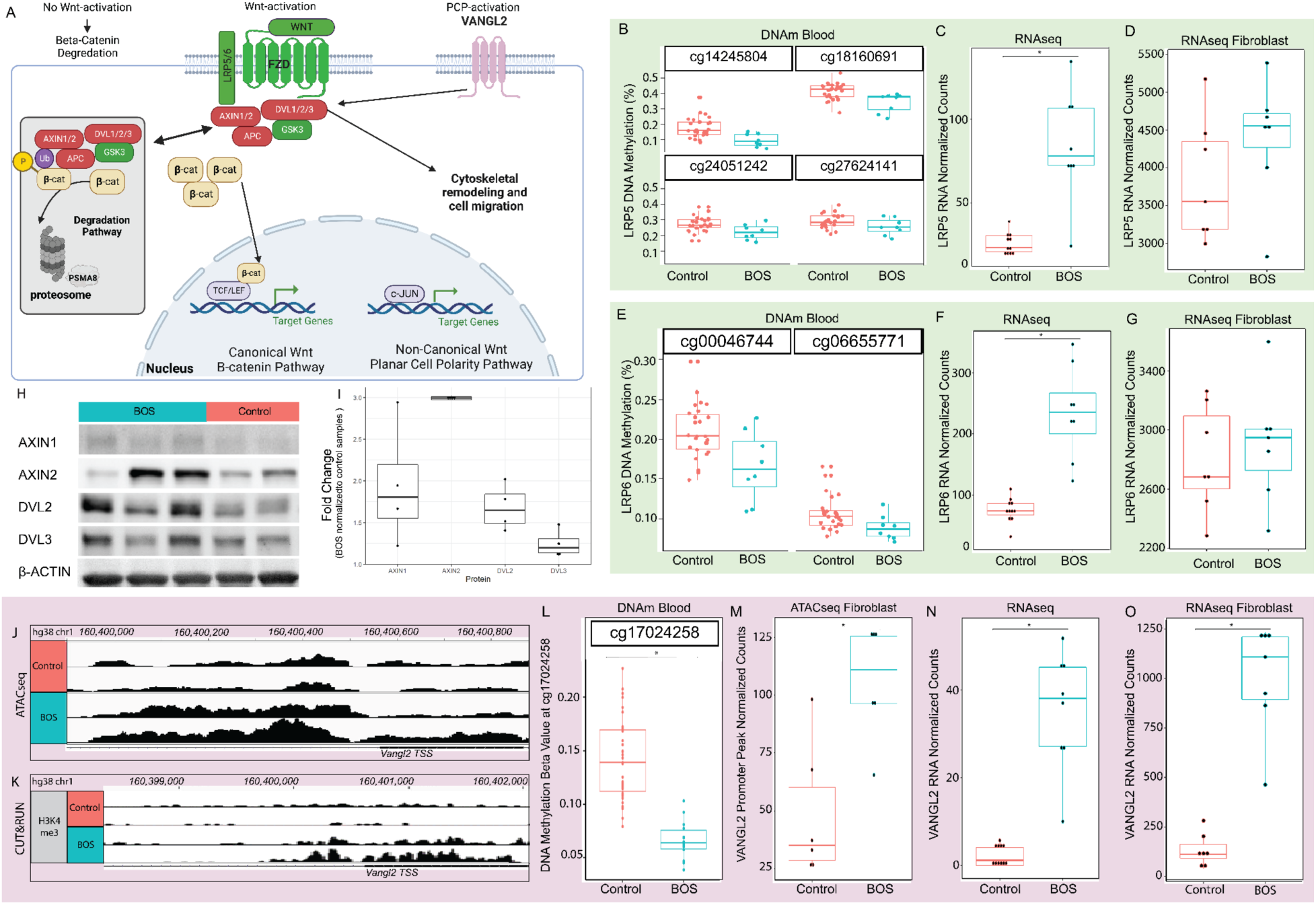
Truncated *ASXL1* dysregulates the canonical and non-canonical Wnt signaling pathways. **(A)** Canonical Wnt signaling pathway (left) is activated when Wnt ligand stimulates its receptors. This inactivates the “β-catenin destruction complex”, allowing nuclear translocation of β-catenin and activation of target genes. Van Gogh-like 2 (VANGL2) intersects with the canonical pathway through activation of DVL to activate non-canonical pathways (right) and cell migration. AXIN1: axis inhibition protein 1; DAAM1: DVL-associated activator of morphogenesis; DVL: disheveled: FZD: frizzled; GSK3β: glycogen synthase kinase 3β; JNK: JUN kinase; RAC: Ras-related C3 botulinum substrate; RHOA: Ras homolog gene family member A; TCF/LEF: T-cell factor/lymphoid enhancer factor. In BOS patient samples, the Wnt pathway co-receptor *LRP5* (green) shows significant **(B)** DNA hypomethylation in blood (Δβ -3.5% to -8.0%) at multiple CpG sites associated with a CpG island, and transcriptional upregulation in **(C)** blood (log_2_FC 1.64) and **(D)** fibroblast RNA-seq. Similarly, *LRP6* (green) shows significant **(E)** DNA hypomethylation in blood (Δβ -2.7% to -4.0%) at multiple CpG sites associated with a CpG island, and transcriptional upregulation in **(F)** blood RNA-seq (log_2_FC 1.63) and **(G)** fibroblast RNA-seq. **(H)** Whole cell lysate (30ug) of BOS and control-derived fibroblasts show Wnt pathway activation at the protein level through staining for AXIN1, AXIN2, DVL2, and DVL3. **(I)** ImageJ quantification showed increases in AXIN2, AXIN1, DVL2, and DVL3. Integration of BOS patient samples showed dysregulation of *VANGL2* across tissue and assay types (pink). In representative patient samples, the *VANGL2* promoter shows **(J)** increased chromatin accessibility and **(K)** increased histone H3K4Me3 marks in BOS compared to control. For BOS samples, *VANGL2* **(L)** is hypomethylated (Δβ -7.6%) at CpG site cg17024258, **(M)** has increased chromatin accessibility at the 5’ UTR (log_2_FC 1.20), and has significant transcriptional upregulation in **(N)** blood RNA-seq (log_2_FC 3.80) and **(O)** fibroblast RNA-seq (log_2_FC 2.55)

We also identified upregulation of *AXIN2* (log_2_FC 0.95) and downregulation of *GSK3β* (log_2_FC -0.28), consistent with canonical Wnt signaling activation. Endogenous *AXIN2* mRNA and protein expression is directly induced by Wnt pathway activation (58, 59) and has been used as a proxy reporter for Wnt pathway signaling. Furthermore, in BOS patient samples, the transcription factors in the Wnt pathway, *TCF7* (log_2_FC 0.81) and *LEF1* (log_2_FC 0.86) are both significantly transcriptionally upregulated.

Western blot quantification of BOS fibroblasts and control fibroblasts using a commercial Wnt antibody kit identified a 3 fold increase in AXIN2, 2 fold increase in AXIN1, 1.5 fold increase in DVL2, and a 1.2 fold increase in DVL3 in BOS patient-derived fibroblasts (**Figure 5H and 5I**). Collectively, our epigenomic, transcriptional and protein data of the Wnt signaling pathways corroborate a distinct and consistent activation of canonical Wnt signaling in BOS samples.

### Multiomics identifies altered epigenetic profile in *VANGL2*, a non-canonical Wnt signaling gene, in BOS

*VANGL2* first came to our attention as one of the most highly overexpressed DEGs in BOS blood and fibroblast RNA-seq data (**Figure 2F, Supplemental Figures 13A and 13B**). Integration of RNA-seq, ATAC-seq and DNAm data again identified *VANGL2* as differentially regulated in all datasets (**Figure 3C and 4B**). In BOS and control fibroblasts, we visualized the ATAC-seq data in IGV which identified a clear increase in reads for BOS patients at the *VANGL2* transcriptional start site (TSS), representing more open chromatin **(Figure 5J**). Similarly, the CUT&RUN results for H3K4me3, an transcription-activating histone mark, showed significantly stronger H3K4me3 binding at the *VANGL2* TSS, for BOS compared to controls (**Figure 5K**). *ASXL1* has been previously shown to affect histone modifications, specifically H3K4me3, H3K27me3 and H2AK119ub, and our data further corroborates that truncating mutations in *ASXL1* result in loss of its gene repressive functions.

In BOS blood samples, we observe the same *VANGL2* transcriptional upregulation coupled with hypomethylation (Δβ= -7.6%) at the TSS of *VANGL2* CpG site cg17024258 (**Figure 5L**) and an inverse relationship between hypermethylation and transcriptional upregulation in individual samples (**Supplemental Figure 13C**). The 5’ UTR of *VANGL2* shows a 2.3-fold increase in chromatin accessibility by ATAC-seq (**Figure 5M**).

These findings are supported by differential gene expression of *VANGL2* in BOS patients versus control samples. As shown in **Figure 2F** with cross-tissue RNA-seq DEG integration, *VANGL2* was significantly overexpressed in both BOS patient blood compared to controls with a normalized RNA count log_2_FC of 3.8 (**Figure 5N**) and in fibroblasts with a log_2_FC of 2.55 (**Figure 5O**).

## DISCUSSION

We utilize a multi-omics approach to profile primary BOS patient samples. For the first time, we directly link *ASXL1* mutations to activation of the canonical and the non-canonical Wnt-signaling pathway, namely the planar cell polarity (PCP) pathway. Both pathways are essential in developmental patterning. The PCP pathway defines cell polarity and is essential to morphogenesis and diverse cellular processes (60). The data demonstrate that upregulation of Wnt pathways in BOS samples can be attributed to disruption of the epigenetic landscape ultimately resulting in transcriptomic dysregulation.

Most of the studies on human ASXL1 function are in the context of myeloid leukemia and propose that truncating *ASXL1* mutations result in haploinsufficiency(11, 36), dominant negative, or gain-of-function effects(31, 37, 61). Our findings (Figures 1D, 2A, 2B, Supplemental Figure 2.) show no evidence of haploinsufficiency or nonsense mediated decay which are consistent with iPSC studies conducted by Matheus et al. (9). We find no significant difference in levels of *ASXL1* transcripts between BOS and control cells, and comparable levels of wild type and pathogenic transcripts in BOS patients. This adds further evidence to existing literature that nonsense mediated decay and haploinsufficiency do not play a role in BOS pathogenesis (37, 62) which is, instead, mediated through mis-expression and impact of a truncated ASXL1 protein.

### High-penetrance mutations in severe monogenic disorders supersede effects of age, sex and genetic background in molecular assays

Although there is strong tissue-specific gene expression(63), we propose that the high-penetrance effect of truncating *ASXL1* mutations results in a set of dysregulated genes across tissues. The 25 common DEGs across BOS blood and fibroblast data support the overarching hypothesis that truncating *ASXL1* mutations exert a strong tissue-independent effect across tissues that have *ASXL1* expression. The functions of these key genes likely drive BOS pathophysiology and targeting these genes may provide a systemic treatment for BOS.

Some key challenges to the study of rare disease are the heterogenity of ages at sample collection and diverse genetic background of patients. Controlling for age-specific effects and genetic background are important but would require statistical power that is near impossible with fewer than 100 patients reported worldwide. The control individuals for DNA methylation were matched for age- and sex-specific effects, while ATAC-seq and RNA-seq controls were matched for sex and genetic background effects. Integration between ATAC-seq, RNA-seq and DNA methylation data, and the identification of common dysregulated genes suggests that the role of truncating *ASXL1* mutations supersedes the effects of age and genetic background.

Integration of ATAC-seq and RNA-seq identified seven genes that were commonly dysregulated (**Figure 3D**), which we tested for age-specific effects since that was a variable which we were not able to sufficiently match. These seven genes, *APBA2, GREM2, KCNQ5, OTUD7A, PELI2, PRXL2A,* and *VANGL2*, were all tested by Lee et al. (64) through transcriptome analysis to classify human genes based on age-specific differential expression analysis and none showed an age-specific signature. These seven genes have important functions in body patterning and organogenesis, nervous system development and ubiquitination regulation. While this does not rule out age-specific differences in potential interacting proteins, it suggests that truncating *ASXL1* effects are not significantly impacted by age.

### *ASXL1* represses canonical and non-canonical Wnt Signaling in normal development

One of the major findings in this study is the link of *ASXL1* mutations and the activation of the canonical and non-canonical Wnt-signaling pathways. The canonical Wnt signaling pathway plays an important role in embryonic development, stem cell maintenance, and differentiation of cells in adults and is highly conserved across species (65). The Wnt signaling pathway is under complex regulation and functions in context-specific manners, dependent on the receptors present on the cell membrane and Wnt ligand-receptor interactions at the time of pathway activation (66).

The molecular mechanisms and specific developmental stages of the interactions between the canonical and non-canonical Wnt pathways are not well elucidated. In the absence of Wnt signaling, β-catenin is degraded by the 26S proteasome, which is composed of two subcomplexes, the 20S proteasome and the regulatory particle(67), (68) (**Figure 5A**). PSMA8 is a subunit of the 20S proteasome, and hypermethylation and transcriptional downregulation of *PSMA8* in BOS patient samples (**Figure 4C and 4D**), suggests downregulation of the 26S proteasome-mediated degradation of β-catenin in BOS patient tissues(67). This supports our findings of aberrant Wnt activation in RNA-seq (**Figure 2E**) and DNA methylation studies (**Figure 4B**) of BOS.

In the Wnt-active state, the “β-catenin destruction complex” is dissociated and culminates in nuclear translocation of β-catenin (42–44) (**Figure 5A**). This nuclear β-catenin binds to transcription factors of the T-cell factor (TCF) and lymphoid enhancer-binding factor (LEF) families, and initiates transcription of Wnt pathway targets (69, 70). *TCF7*, upregulated in BOS samples, is an important regulator of self-renewal and differentiation in multipotential hematopoietic cell lines (71), which could lend insight to the pathogenesis of myeloid cancers with truncating *ASXL1* mutations.

In contrast, the non-canonical pathways, including the PCP pathway which defines cell polarity and migration, are characterized by their β-catenin-independent regulation of crucial events during embryonic development (72). However, experiments that indicate their binary utilization may be model-system and context-specific (47). Given the complex interplay and cross-talk at almost every level of the Wnt signaling pathways that we are just beginning to unravel, van Amerongen & Nusse suggest moving towards a more integrated view rather than the outdated binary classification (47).

### Dysregulation of the Wnt signaling pathway identified in BOS samples may explain phenotypic presentations in BOS patients

Wnt signaling plays a key role in hair growth and development of hair follicles. In particular, *WNT10B*, which is transcriptionally upregulated in BOS samples, promotes differentiation of primary skin epithelial cells towards the hair shaft in mice and elongation of the hair shaft in isolated rabbit whisker hair follicles (73–75). Intriguingly, many providers and parents have noted BOS patient phenotype of hypertrichosis with rapidly growing hair and nails (25). This lends support to our hypothesis that activation of Wnt signaling may play an important role in BOS patient pathophysiology.

Other significant Wnt signaling DEGs in our datasets include *WNT1*, important in regulating cell fate and patterning during embryogenesis, and *WNT7A*, a key embryonic dorsal vs ventral patterning gene and a key regulator of normal neural stem cell renewal, proliferation, and differentiation. *WNT7A* activates the PCP pathway and *Wnt7a* overexpression has been shown to induce *Vangl2* overexpression and impair neurulation in mouse neural stem cells through aberrant *Vangl2* polarized distribution (76, 77)

### Upregulation of non-canonical Wnt signaling gene *VANGL2* in BOS drives clinical phenotypes

*VANGL2* was one of the most highly dysregulated genes in all our datasets with a 9-fold increase in gene expression occurring via epigenetic dysregulation (**Figures 5J - 5O**). *VANGL2* is thought to regulate Wnt protein distribution (78) and in its non-canonical role has been shown to play critical roles in neurulation, cardiac development, kidney-branching morphogenesis and regulation of hematopoiesis (48, 79–81). The PCP pathway is a highly conserved non-canonical Wnt signaling pathway important in establishing and maintaining polarity during morphogenesis. The asymmetric localization on the plasma membrane of VANGL2 is needed for signal transduction, and subsequent polarization and organization of cells (82). The PCP pathway is also crucial for neural tube closure (83) and it has been suggested that the gradient of Wnt activity helps establish VANGL2 polarity in the neural plate during neurulation (84), with canonical Wnt signaling required for neural crest induction, and non-canonical Wnt pathway required for neural crest migration (85).

Knockout and loss-of-function studies in *Vangl2* identified significant reduction in spine density and dendritic branching in primary culture rat hippocampal neurons (48). In humans, homozygous mutations in *VANGL2* are embryonic lethal and cause craniorachischisis, a very severe neural tube defect encompassing anencephaly and bony defects of the spine, in mice (86) whereas heterozygous *VANGL2* mutations are embryonic lethal and detected in miscarried human fetuses with severe cranial neural-tube defects (87). Neural crest cells contribute to nervous system and craniofacial development, and are critical for cardiac outflow septation and alignment (88). Perhaps not surprisingly, *Vangl2* knockout mice exhibit cardiac outflow tract malformations and septal defects (79), as do BOS patients (25). These findings suggest that the severe neural phenotype, distinctive craniofacial features, and cardiac defects of patients with BOS (25) may be due, at least in part, to the dysregulation of *VANGL2* and downstream effects on neural crest migration.

Although the interplay between the canonical Wnt and non-canonical Wnt pathways is not well understood, recent literature has suggested that these pathways exert reciprocal pathway inhibition through competition for a common downstream coreceptor, Frizzled (*FZD*) (72, 89), most evident during tissue regeneration and development (90, 91). The asymmetric localization of *FZD3* required for proper function is dependent on anchoring by *VANGL2* through physical interaction of the two proteins (92). We identified upregulation of *FZD3*, which regulates establishment of the non-canonical Wnt / PCP pathway and is involved in neural crest cell migration and neural tube closure (93, 94). These results are supported by molecular characterization of truncating *ASXL1* mutations conducted in iPSCs (9).

A limitation of this study is that BOS pathophysiology is shaped by cellular context and developmental stage and our study is focused on differentiated cell types. Cellular and developmental context can drastically affect the molecular effects of epigenetic modifiers such as *ASXL1*. The effects of *ASXL1* begin in early embryogenesis (95) so embryonic stem cells would be the ideal model to recapture cell types where *ASXL1* is maximally impacting progenitors of clinical phenotypes. These multi-omics approaches will benefit from being performed in patient-derived stem-cell models and through samples generated with introduction of truncating *ASXL1* mutations using DNA-editing systems. Moving into stem-cell models will yield additional insights into the role of *ASXL1* in stem cell homeostasis and differentiation.

In conclusion, we present a comprehensive multi-omics analysis of BOS using primary patient samples. This study encompasses the largest omics study of patients with BOS. We present the first analyses of RNA-seq and ATAC-seq conducted on primary BOS patient samples. Through DNA methylation assays, identifying binding proteins from CUT&RUN sequencing, deciphering chromatin accessibility through ATAC-seq, and gene expression from RNA-seq, we add to growing literature on *ASXL1* and the pathogenesis of truncating mutations. Our integrated methods identified dysregulation of key transcripts such as *VANGL2* through direct chromatin modifications. This suggests that physiological levels of truncating *ASXL1* mutation lead to loss of the repressive function of *ASXL1*. Importantly, pathways that are dysregulated across multiple assays are involved in neural development, which shed light on the severe intellectual and neural features in BOS patients, and Wnt/β-catenin and non-canonical Wnt PCP pathways which represent key, drug-targetable pathways. Our findings have major implications for identifying potential therapeutics for diseases harboring *ASXL1* mutations.

## METHODS

### Statistics

RNA-seq, ATAC-seq, and CUT&RUN reads were aligned to the human genome (hg38) and featureCounts (v1.6.5) was used to generate count matrices for genes or chromatin regions. DESeq2 (v1.24.0), which internally normalizes sample library size and utilizes the negative binomial distribution for testing, was used to identify differentially expressed genes (RNAseq) or differentially accessible chromatin regions (ATAC-seq, CUT&RUN) after adjusting for sample sex, which was identified as a covariate to adjust for through PCAs generated based on the top 500 most variably expressed genes. Genes or regions were identified as significant if Benjamini-Hochberg adjusted p-values (p_adj_) were less than 0.05. We further filtered for highly dysregulated significant genes or regions using an abs(log_2_FC) ≥ 0.58, corresponding to an absolute fold change ≥ 1.5 with reference to control samples. Gene ontology over-enrichment tests were completed using clusterProfiler v3.12.0 by submitting differentially expressed genes against all genes from the Gencode hg38 annotation, version 31. Gene ontologies were classified as significantly enriched when p-adjusted (Benjamini-Hochberg) was less than 0.05 (hypergeometric test).

DNA methylation beta (β) values were calculated using minfi(96), and differentially methylated sites were identified by running Limma regression modeling(97), while adjusting for the covariates of age, sex, and ethnicity. Our significance cut-offs were FDR < 0.05 (Benjamini-Hochberg). We filtered for significantly methylated CpG sites with abs(Δβ) > 0.10, where Δβ represents the difference in average DNAm (β) between BOS and controls. GREAT (56) was run on significant CpG sites to identify biological mechanisms that were dysregulated, filtering for gene ontologies with p_adj_ < 0.05 for significance.

### Study Approval

This project was approved by the UCLA IRB# 11-001087. In conjunction with Dr. Bianca Russell and the ASXL Biobank at UCLA, skin punch biopsies and blood samples from ASXL patients and parents were collected, as well as deep phenotyping and review of medical records. We used primary sample types to examine the direct role of the mutation on patient tissue. We used controls with similar genetic backgrounds to control for baseline genetic effects. All individual level-data was de-identified prior to analysis and samples were collected for follow-up experiments. We collected blood from 12 BOS individuals and 38 controls, and skin punch biopsies from 6 individuals and 13 controls. We used two independent samples from one individual to filter out noise of genetic background. Whole blood was processed to isolate DNA, RNA and PBMCs and skin-punch biopsy samples were processed to create patient-derived skin fibroblasts in the Translational Pathology Core Laboratory at UCLA.

### Cell Culture

Patient derived fibroblast cell lines were grown in DMEM (Gibco™, #11-995-073), 10% FBS, 1% Non-essential Amino Acid (100X, Gibco™, 11140-050) and 1% PenStrep at 37℃ in 5% CO_2_ incubators. For transfection experiments, HEK293T cells and CACO2 cells were grown in DMEM (Gibco™, #11-995-073), 10% FBS, and 1% PenStrep at 37℃ in 5% CO_2_ incubators. All cell lines were tested for mycoplasma (MycoAlert PLUS Mycoplasma Detection Kit, Lonza #LT07-318) on a monthly basis.

### Western Blots

Whole cell lysate was extracted with Cell Lysis Buffer (10X, Cell Signaling #9803S), cytoplasmic and nuclear extractions using the Thermo Scientific CE/NER Kit (#78833), and histones were extracted as per established protocol (98). 1X Halt™ Protease and Phosphatase Inhibitor Cocktail (100X, Thermo Scientific™ #PI78442) was used. Protein was quantified using the Pierce BCA Protein Assay Kit (Thermo Scientific #23225). Samples were resolved by SDS polyacrylamide gel electrophoresis on 4%-15% Criterion^TM^ TGX Stain-Free^TM^ Protein Gel (BioRad #5678083) using 30μg of whole cell lysate, 10μg of cytoplasmic extract, 3ug of nuclear extract or 1ug of acid-extracted histone per well for 30 minutes at 190V. Proteins were then transferred to 0.2uM nitrocellulose membranes (BioRad #1704271) using the semi-dry TransBlot for 7 min at 25 mA. The membranes were blocked with 5% milk in 1x TBST and then probed with primary antibodies (1:100) overnight at 4℃, and with secondary antibodies at room temperature for one hour according to manufacturer dilutions. Blots were imaged on a Biorad ChemiDoc. For a full list of antibodies, see **Supplemental Table 6**.

### Blood Sample Collection and Processing

Peripheral blood samples were collected into EDTA tubes for DNA extraction and PBMC extraction, CPT tubes for PBMC extraction and PAXgene™ Blood RNA Tube (BDBiosciences, 762165) for RNA extraction. Whole blood DNA was extracted using the Qiagen Blood and Cell Culture DNA Midi Kit (Qiagen #13343), and whole blood RNA was extracted using the MagMAX™ for Stabilized Blood Tubes RNA Isolation kit (Thermo Fisher Scientific, #4451893). All samples were quantified on the nanodrop, Qubit 4.0 and Tapestation Bioanalyzer for sample quality control. DNA and RNA were stored at -80C.

### RNA-sequencing

BOS patient-derived dermal fibroblasts, and sex-matched controls were grown to 80-90% confluency in T75 flasks before total RNA extraction via RNeasy Plus Mini Kit (Qiagen, 74136). Whole blood samples were collected and purified using MagMAX for Stabilized Blood Tubes RNA Isolation Kit (Qiagen, 3341894) with DNase treatment. All samples were quantified on the Qubit 4.0 and Tapestation Bioanalyzer 4150 using High Sensitivity RNA Tapestation (Agilent, 5067-5579) to ensure high-quality RNA (RIN >= 9.0 for fibroblasts, RIN <=5.3 for blood were used for subsequent library construction). Blood samples were first treated with QIAseq FastSelect −Globin (Qiagen, 334376). All samples were then prepared with QIAseq FastSelect −rRNA HMR (Qiagen, 334386) and lllumina Truseq® Stranded Total RNA Library Prep Gold (Illumina, 20020599). All libraries were quantified on the Qubit 4.0 and library quality was assessed using Agilent D1000 DNA Tapestation (Agilent, 5067-5582) before multiplexing and sequencing on a NovaSeq flow cell for a minimum of 30 million paired-end 150bp reads per 17sample. *Fastq* files were processed through our best practice bioinformatic in-house pipeline (**Supplemental Methods and Supplemental Figure 16**). Multidimensional scaling based on the top 500 genes with the highest variance was performed and verified that BOS samples clustered together and away from controls (**Supplemental Figures 14A and 15A**). We confirmed effective rRNA removal (**Supplemental Figures 14B and 15B**). Differential expression analysis was performed, adjusting for sample sex, and genes were identified as significant at p_adj_ < 0.05 (Benjamini-Hochberg). We further filtered our differentially expressed genes (DEGs) for abs(log_2_FC) ≥ 0.58, corresponding to an absolute log fold change ≥ 1.5.

### ATAC-Seq

Patient-derived fibroblast lines were cultured *in vitro* to 80-90% confluency in T75 flasks and 50,000 freshly isolated cells per line were treated with Tn5 transposase as per established protocol (99). After purification and PCR amplification for library generation, the libraries were double-sided bead purified using AMPure XP beads (Beckman, A63881) to remove contaminating primer dimers. All libraries were quantified on the Qubit 4.0 and library quality was assessed using Agilent High Sensitivity DNA Tapestation (Agilent, 5067-5584) before multiplexing and sequencing on a Nextseq 550 for a minimum of 40 million paired-end reads per sample with 75bp length. *Fastq* files were processed through our best practise bioinformatic in-house pipeline (**Supplemental Methods and Supplemental Figure 17**). Technical replicates clustered closely on PC analysis and were collapsed during analysis.

### CUT&RUN

CUT&RUN (Cleavage Under Targets & Release Using Nuclease) libraries were prepared with the established CUT&RUN Assay Kit (Cell Signaling, #86652S). The following antibodies were used: IgG isotype (Cell Signaling, #66362, negative control, 1:20), H3K4me3 (Cell Signaling, #9751S, 1:50) and H3K27me3 (Cell Signaling, #9733S, 1:50) according to protocol. For DNA purification, the DNA Purification Buffers and Spin Columns for ChIP and CUT&RUN (Cell Signaling, #14209), with elution in 50 ul.

CUT&RUN libraries were prepared using the SimpleChIP ChIP-seq DNA Library Prep Kit for Illumina (Cell Signaling, #56795S) with SimpleChIP ChIP-seq Multiplex Oligos for Illumina (Cell Signaling, #47538S) according to manufacturer guidelines, with modifications for CUT&RUN as noted in the CUT&RUN Assay Kit (Cell Signaling, #86652S). All libraries were quantified on the Qubit 4.0 and library quality assessed using Agilent High Sensitivity DNA Tapestation (Agilent, 5067-5584) before multiplexing and sequencing on a Nextseq 550 Mid Output Kit (150 cycles) with 1% PhiX for a minimum of 5 million paired-end reads per sample with 50bp read length. Samples were analyzed using the established NF-Core CUT&RUN analysis pipeline(100).

### DNA methylation

We used DNA methylation generated from BOS and control blood and fibroblasts and processed as described in (49). We calculated the absolute difference between the means of the β value for BOS patients versus controls for each CpG to obtain the delta beta (Δβ) value. We filtered for highly differentially methylated sites (|delta beta (Δβ)| > 5%). Significant CpG sites were identified if they were below FDR < 0.05. To identify biological mechanisms that are dysregulated, we queried these CpG sites using GREAT (56). We filtered for gene ontologies with p_adj_ < 0.05 for significance.

### RT-qPCR

RT-qPCR was conducted using TaqPath™ 1-Step RT-qPCR Master Mix, CG (Applied Biosystems, A15300) according to protocol (Document #100020171, Rev 1.00) on QuantStudio™ 3. 10ng of RNA was used per well in 20ul total well volume, and each sample and primer combination was conducted in triplicate. Machine settings were for Standard Curve and Fast run mode. RNA isolated from patient blood and patient-derived fibroblast cell lines were subject to RT-qPCR to validate RNA-seq findings. RNA isolated from transfection experiments were subject to RT-qPCR to identify effects of the ASXL1 truncated plasmids or ASXL1 siRNAs on cell transcriptome.

## Public Data Sets

https://portal.gdc.cancer.gov/projects/TCGA-LAML.

## Author Contributions

VA, ZA and IL designed and conceptualized the study and wrote the paper. IL, MS and AW performed data generation and transcriptomic and epigenomic data analysis for patient-derived samples. BR coordinated sample collection. ZA and RW performed data generation and analysis of DNA methylation data. All authors contributed to the writing of and the editing of the manuscript.

## Acknowledgements

This work was supported by the following funding sources awarded to V.A.A.: NIH DP5OD024579 and the ASXL Research Related Endowment Pilot Grant (2020–2022). IL was supported by NIH T32 GM008042. ZA and RW are supported by the ASXL Research Related Endowment Pilot Grant (2020–2022), the Canadian Institutes of Health Research (CIHR) grants (IGH-155182 and MOP-126054), and the Ontario Brain Institute (province of Ontario Neurodevelopmental Disorders (POND) network (IDS11-02)) grants to RW. We would like to acknowledge the families and patients who participated in this study and donated their time and efforts to make this possible. We would also like to acknowledge Elisabeth McGee for her work in coordinating sample collection and The California Center for Rare Disease for their support and building of critical infrastructure for these studies. Some of the figures in this paper were created with BioRender.com.

## SUPPLEMENTAL MATERIALS

### SUPPLEMENTAL FIGURES

**Supplemental Figure 1:**
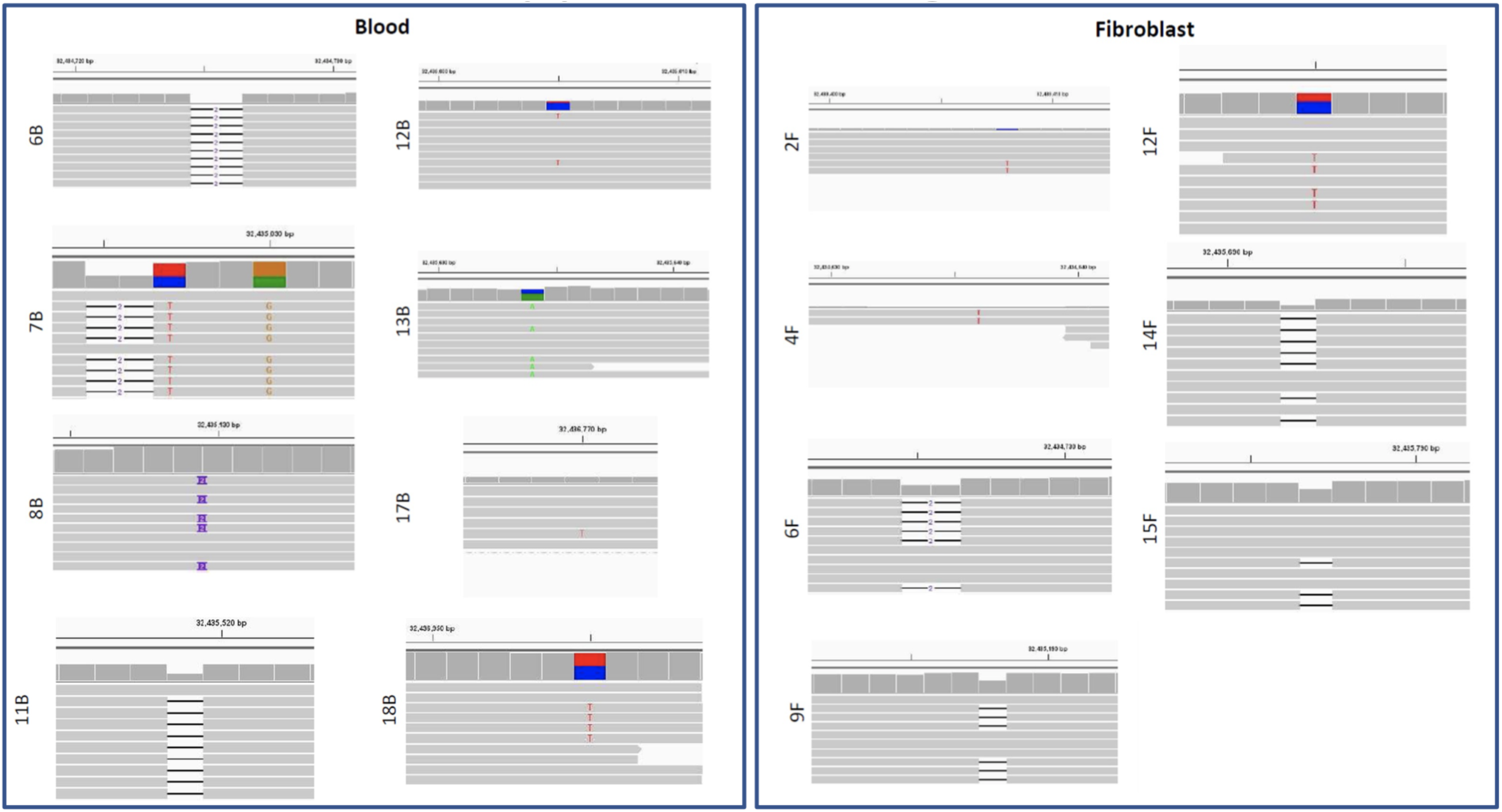
Validation of *ASXL1* mutation and identity in BOS Samples. Representative RNAseq reads for each BOS patient blood and fibroblast sample show patient mutation. Patient ID numbers (Figure 1, Table S1) and sample type (F = fibroblast, B = blood) are listed on the left of each insert. Genomic positions for hg38 *ASXL1* (20q11.21) are shown above each insert. For complete coverage see Table S9.

**Supplemental Figure 2:**
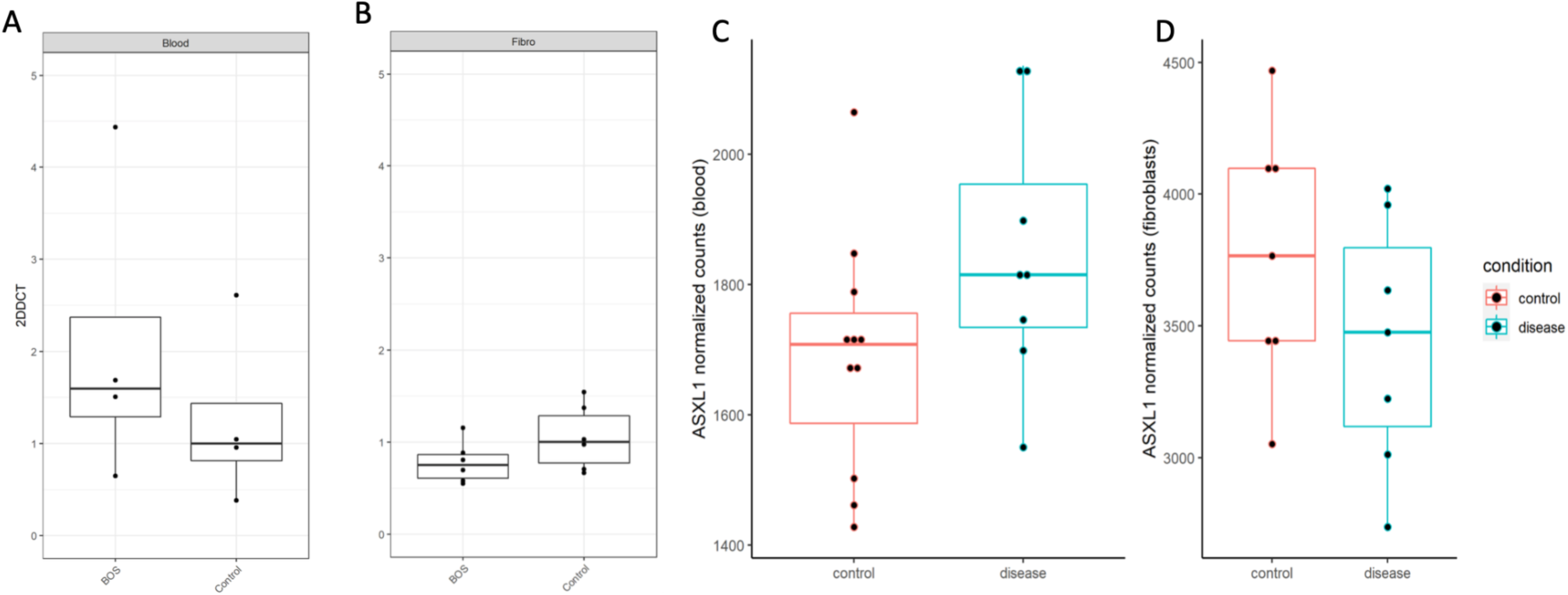
Assessment of *ASXL1* expression in patient and control blood and fibroblast samples. Quantitative real-time polymerase chain reaction (qPCR) relative gene expression calculated by the delta-delta Ct method (2−ΔΔCt) do not show significant difference in *ASXL1* expression in (**A**) blood or (**B**) fibroblasts between representative BOS and control samples. RNA-sequencing DESeq2 normalized counts also do not show significant differential expression of *ASXL1* between BOS and control samples in (**C**) blood (log2FC = 0.11, padj = 0.29) or (**D**) fibroblasts (log2FC = -0.01, padj = 0.95).

**Supplemental Figure 3:**
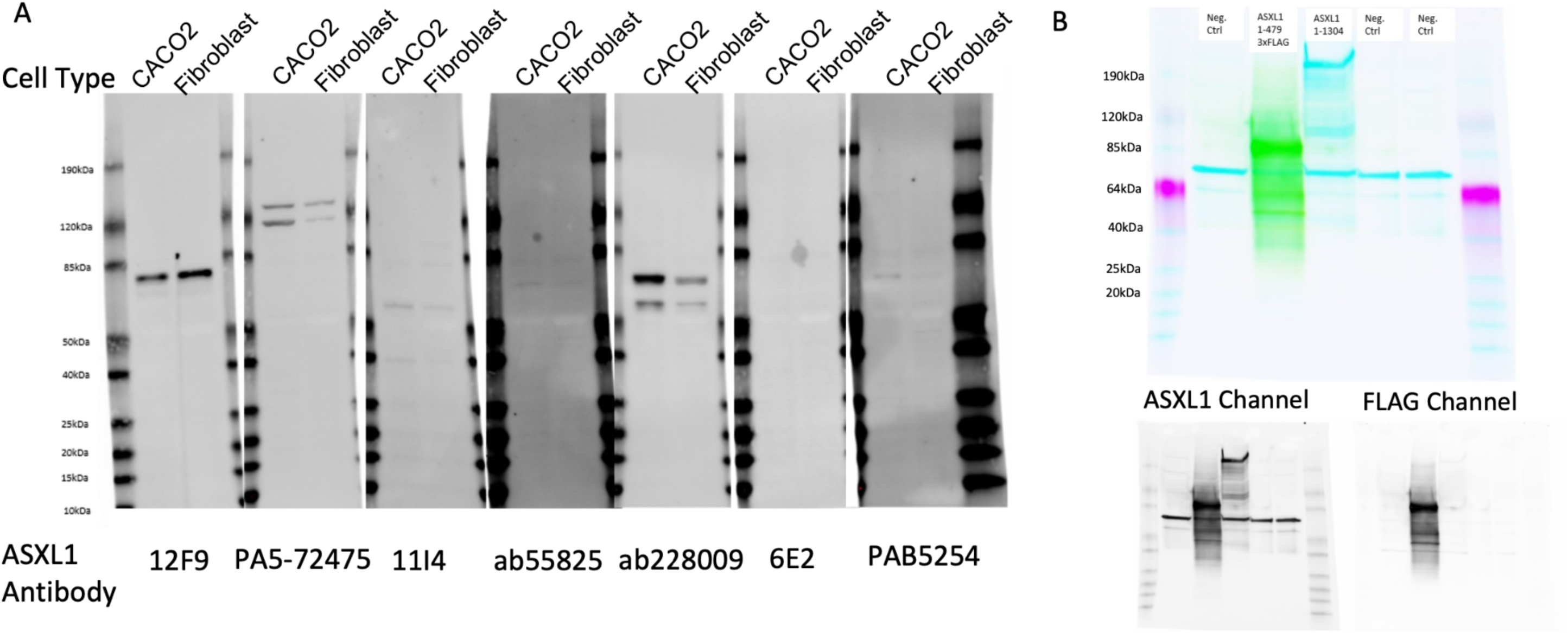
ASXL1 Antibody Testing. (**A**) An array of antibodies targeting *ASXL1* were tested on whole cell lysate (20ug) from CACO2 cells and fibroblast cells. Only antibody #ab228009 identified higher ASXL1 expression in CACO2 cells, which was expected based on ProteinAtlas data (Supplementary Table 4). (**B**) To see whether ASXL1 antibody #ab228009 could identify an exogenous and/or endogenous ASXL1 correctly, we transfected truncated ASXL1 plasmids into HEK293T cells and resolved 10ug nuclear extract on western gel. *ASXL1* antibody ab228009 identified the truncated ASXL1 plasmids: ASXL1 1-479 + FLAGx3, and ASXL1 1-1304. ASXL1 (below, left) and FLAG (below, right) staining for the ASXL1 1-479 + FLAGx3 cell lysate showed complete overlap.

**Supplemental Figure 4:**
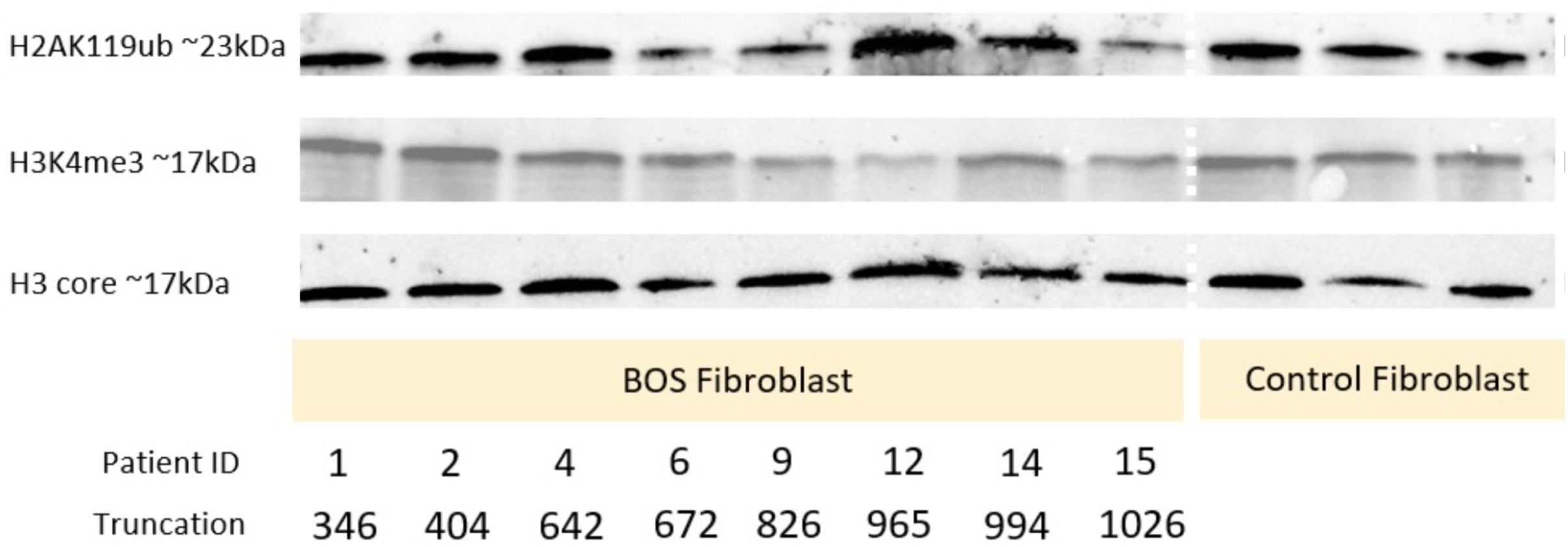
Extended histone H2AK119ub and H3K4me3 western blots. Histone extracts (1ug) from BOS patient and control fibroblast were stained with H2AK119ub and H3K4me3 antibody. No visible or quantifiable difference was identified between patient and control histone marks.

**Supplemental Figure 5:**
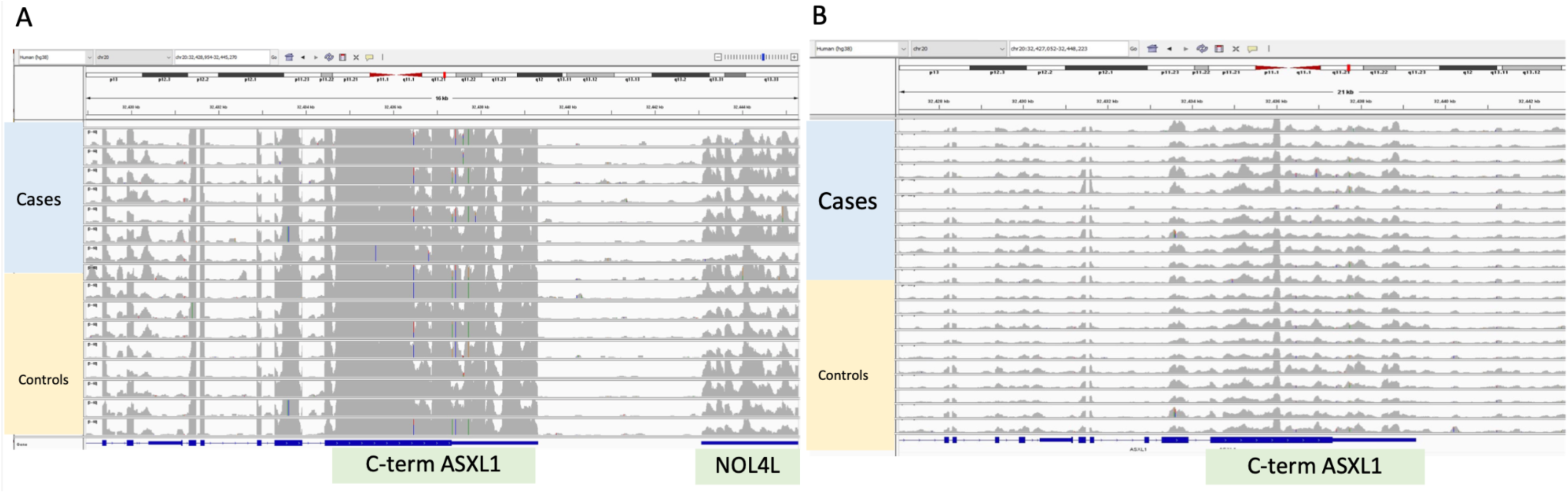
Normalized ASXL1 reads in BOS patients and controls. Integrative Genome Viewer is used to show normalized read counts for ASXL1 in BOS patients and controls in (**A**) fibroblast samples and (**B**) blood samples. Similar normalized levels of ASXL1 are observed for BOS cases and controls in both tissue types at the C-terminus.

**Supplemental Figure 6:**
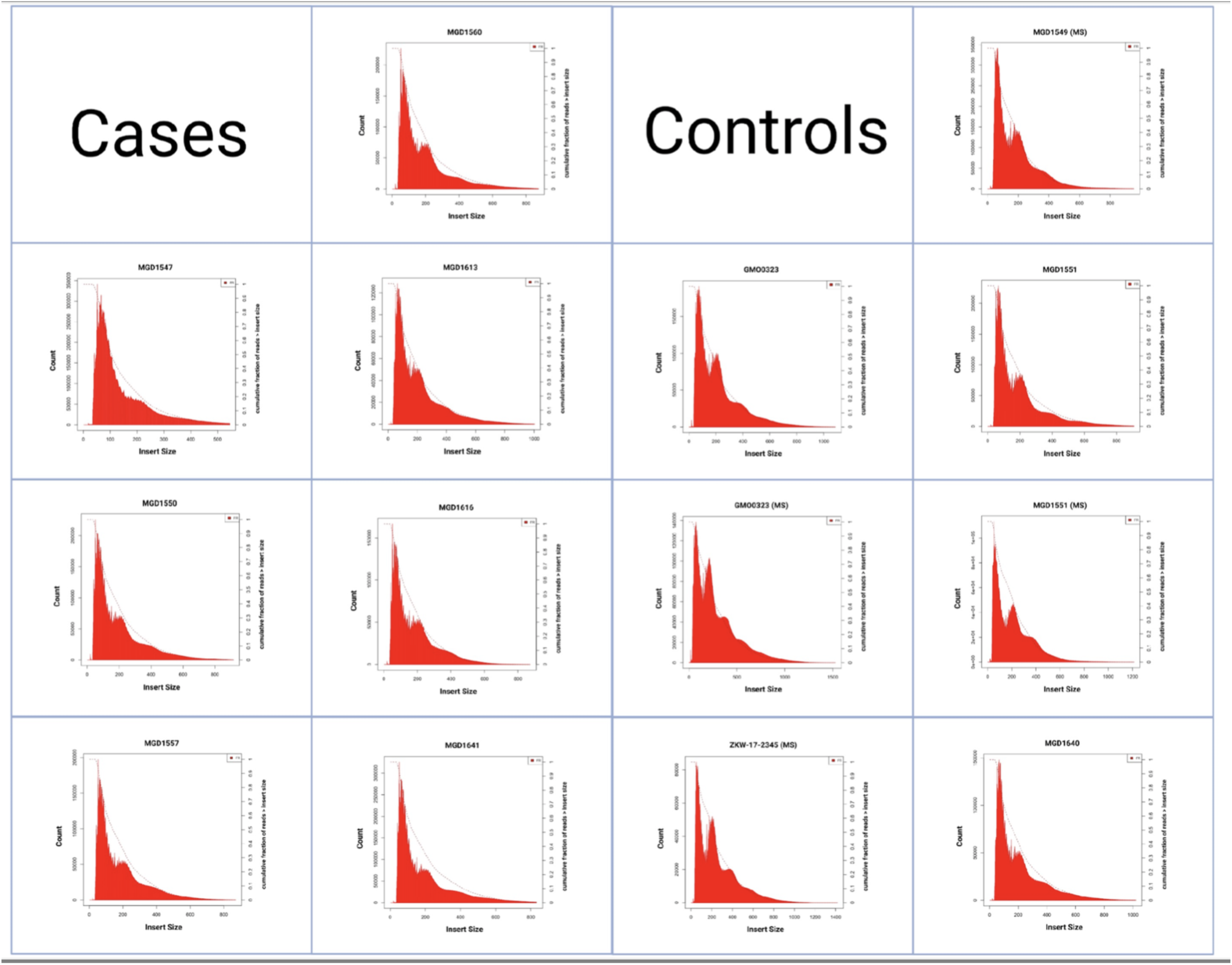
ATAC-seq Sample Fragment Length. Fragment length distribution plots for each ATAC-seq sample are shown. We identify more short fragment lengths in each sample, which correspond to more open chromatin, and decreasing amounts of large fragments, which correspond to less open chromatin.

**Supplemental Figure 7:**
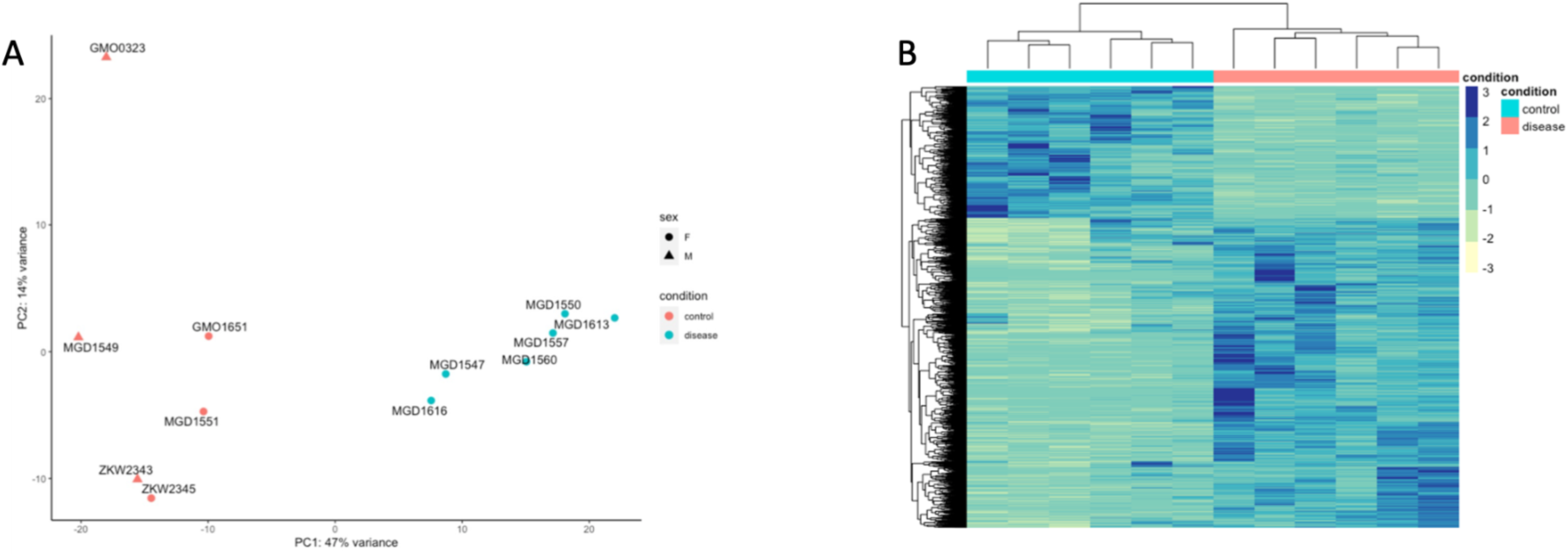
ATACseq Fibroblast Library QC. (**A**) Principal component analysis with adjustment for sex identifies moderate separation of patient and control samples. (**B**) Heat map of ATAC-seq fibroblast samples identifies clear separation of BOS patient and controls. We also see more DEGs corresponding to open chromatin peaks in BOS samples than controls.

**Supplemental Figure 8:**
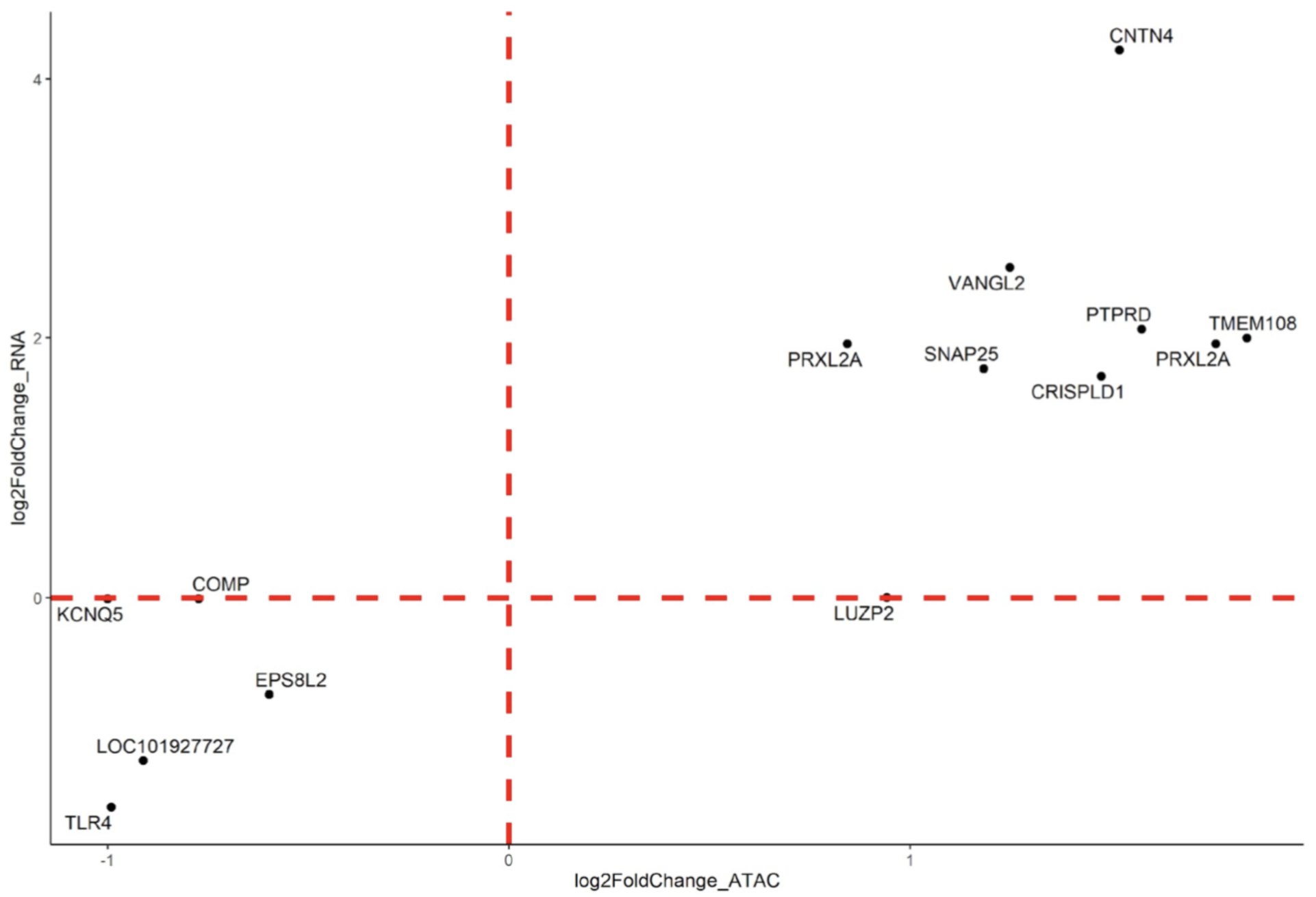
Integration of differential open chromatin at transcriptional start sites with transcriptomic dysregulation in BOS fibroblasts. Significant DEGs from BOS RNAseq fibroblast data was integrated with significant differential chromatin accessibility peaks at the transcriptional start site from BOS ATACseq fibroblast data. DEGs that overlapped are shows with the log_2_FC for RNAseq (y axis) and log_2_FC for ATACseq (x axis).

**Supplemental Figure 9:**
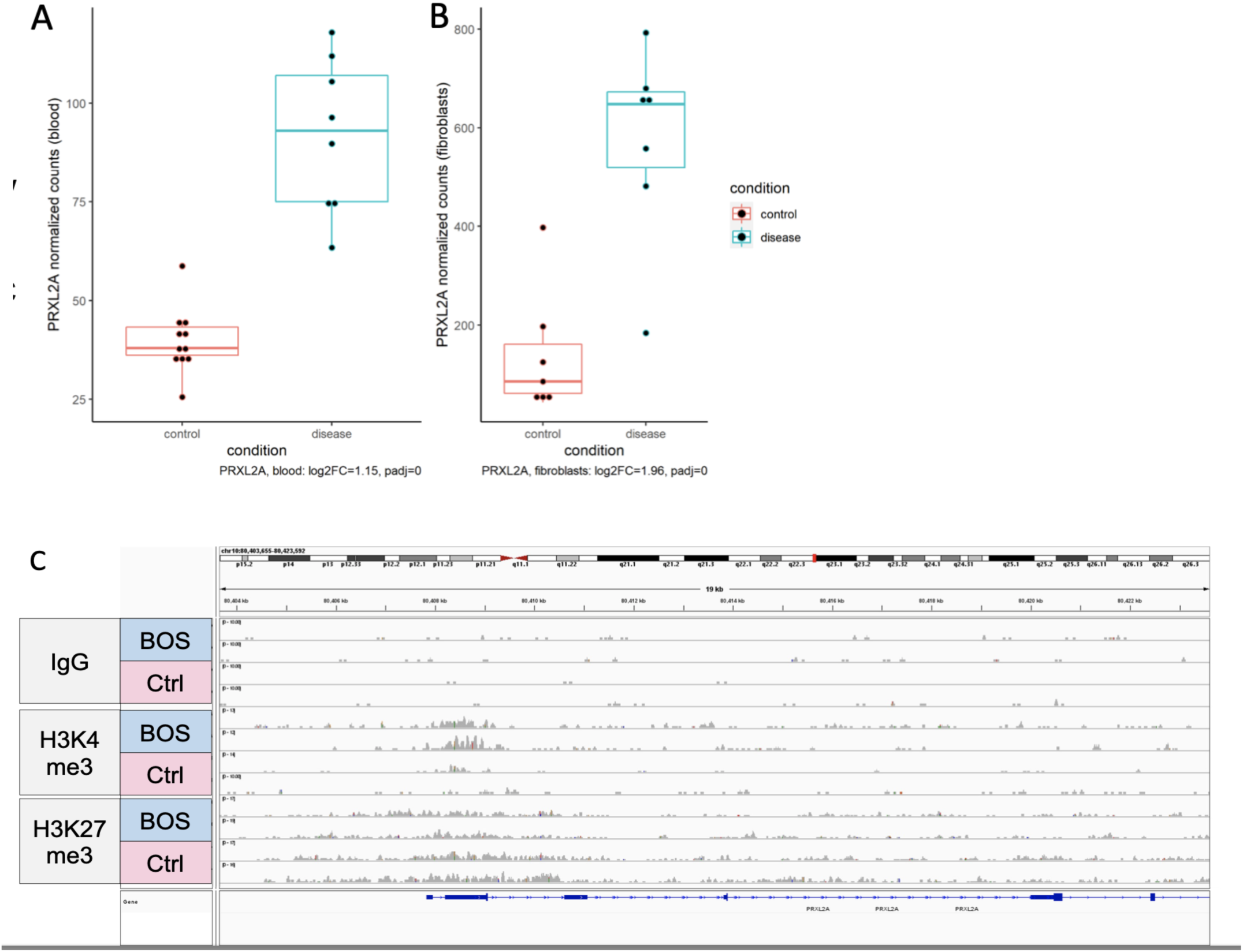
Multi-ome integration identifies *PRXL2A* dysregulation in BOS samples. RNA-sequencing DESeq2 normalized counts show significant differential expression of *PRXL2A* between BOS and control samples in (**A**) blood (log2FC = 1.15, padj = 0) or (**B**) fibroblasts (log2FC = 1.96, padj = 0). (**C**) CUT&RUN identifies increased H3K4me3 binding at the *PRXL2A* transcriptional start site.

**Supplemental Figure 10:**
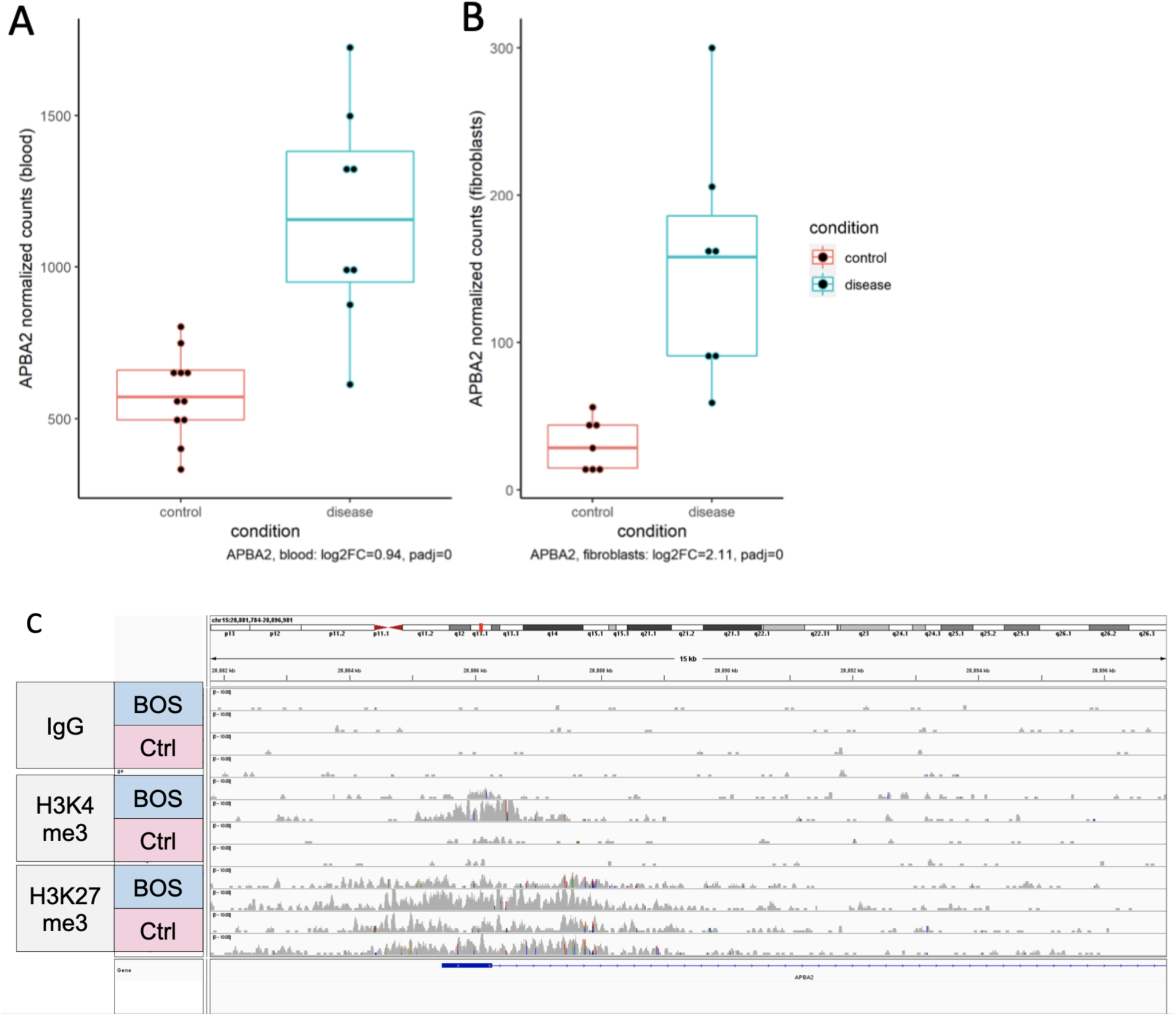
Multi-ome integration identifies *APBA2* dysregulation in BOS samples. RNA-sequencing DESeq2 normalized counts show significant differential expression of *APBA2* between BOS and control samples in (**A**) blood (log2FC = 0.94, padj = 0) or (**B**) fibroblasts (log2FC = 2.11, padj = 0). (**C**) CUT&RUN identifies increased H3K4me3 binding at the *APBA2* transcriptional start site.

**Supplemental Figure 11:**
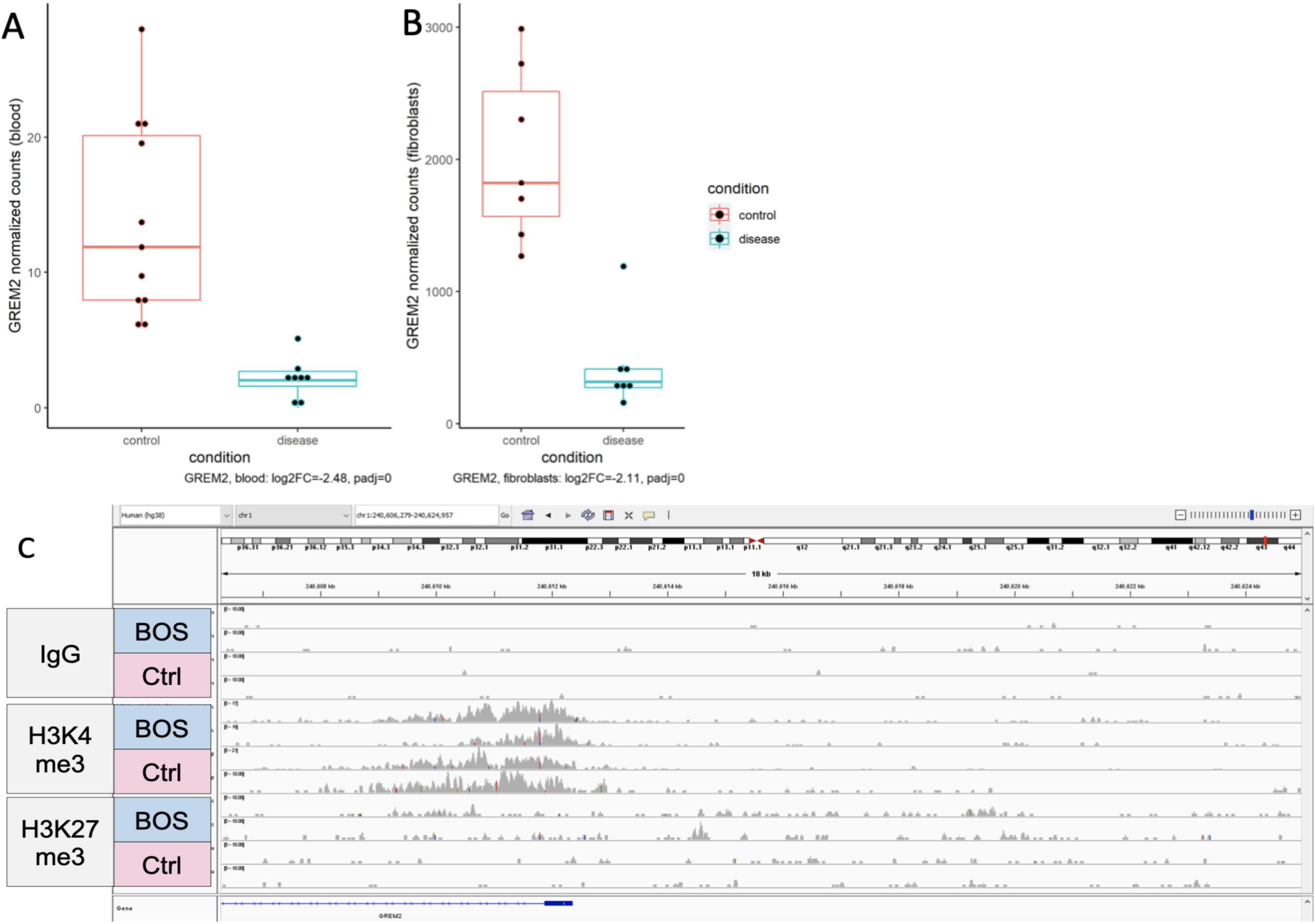
Multi-ome integration identifies *GREM2* dysregulation in BOS samples. RNA-sequencing DESeq2 normalized counts show significant differential expression of *GREM2* between BOS and control samples in (**A**) blood (log2FC = -2.48, padj = 0) or (**B**) fibroblasts (log2FC = -2.11, padj = 0). (**C**) CUT&RUN identifies increased H3K4me3 binding at the *GREM2* transcriptional start site.

**Supplemental Figure 12:**
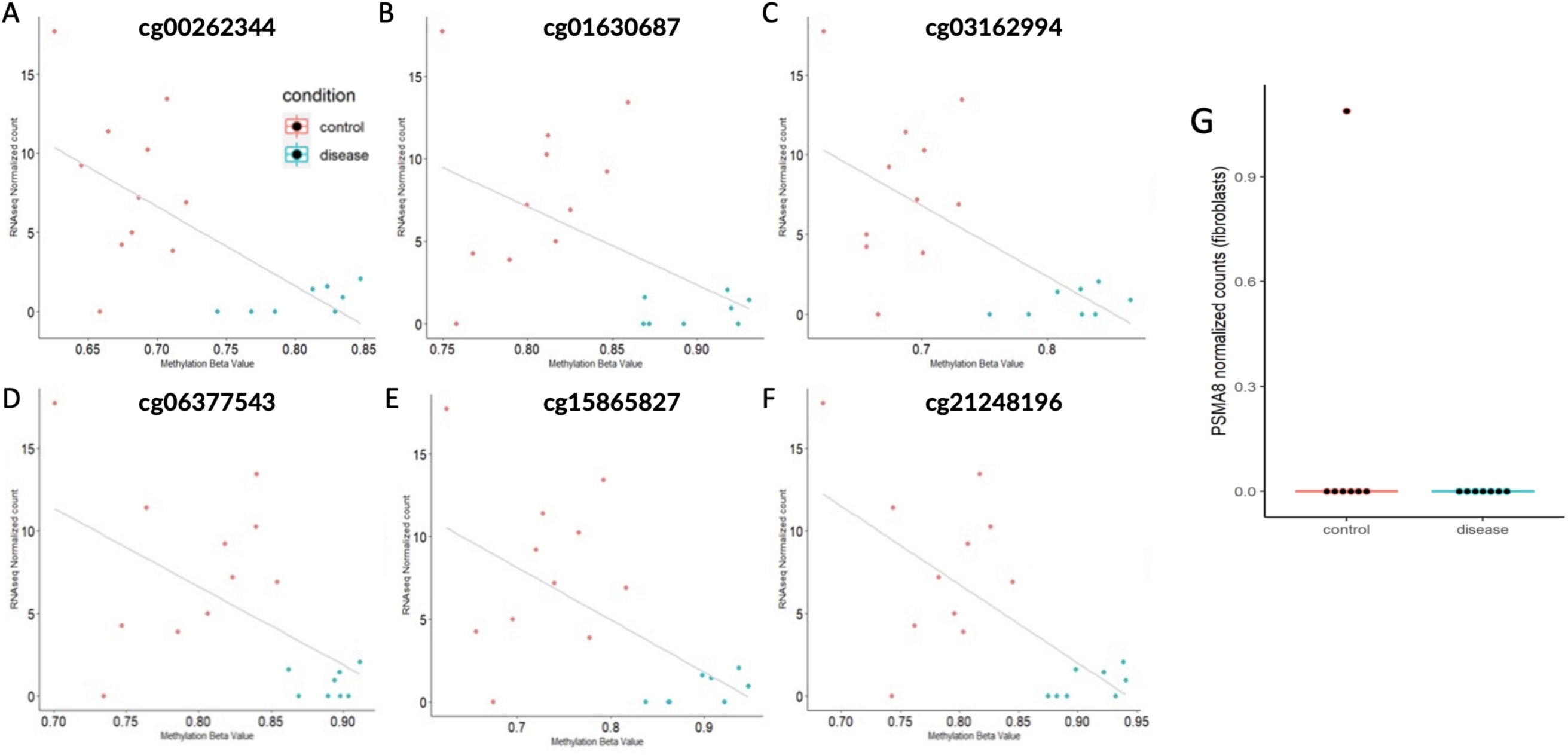
Differentially methylated CpG sites at PSMA8 in BOS patient blood. BOS patient (blue) and control (red) samples transcriptomic normalized values from blood RNAseq (x axis) and methylation beta values from blood DNAm (y axis) were plotted for CpG site (**A**) cg00262344 (**B**) cg01630687 (**C**) cg03162994 (**D**) cg06377543 (**E**) cg15865827 (**F**) cg21248196. At each CpG sites, BOS patient samples were hypermethylated and transcriptionally downregulated compared to controls. (**G**) RNAseq did not identify any *PSMA8* reads in fibroblast samples.

**Supplemental Figure 13:**
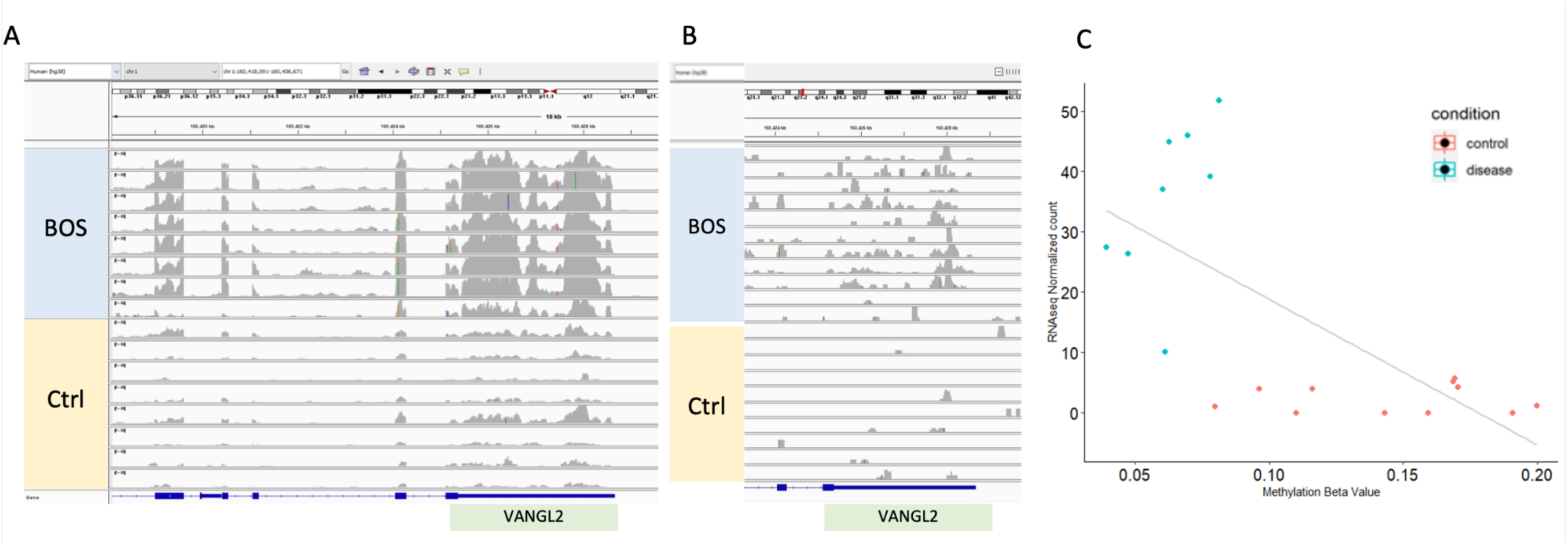
VANGL2 is one of the most highly overexpressed transcripts in BOS patient samples. Normalized *VANGL2* reads in BOS patients and controls (**A**) fibroblast samples and (**B**) blood samples show increased expression in BOS samples. (**C**) At CpG site cg17024258 located at the *VANGL2* transcriptional start site, BOS patient samples are hypomethylated (Δβ -7.6%) and transcriptionally upregulated compared to controls.

**Supplemental Figure 14:**
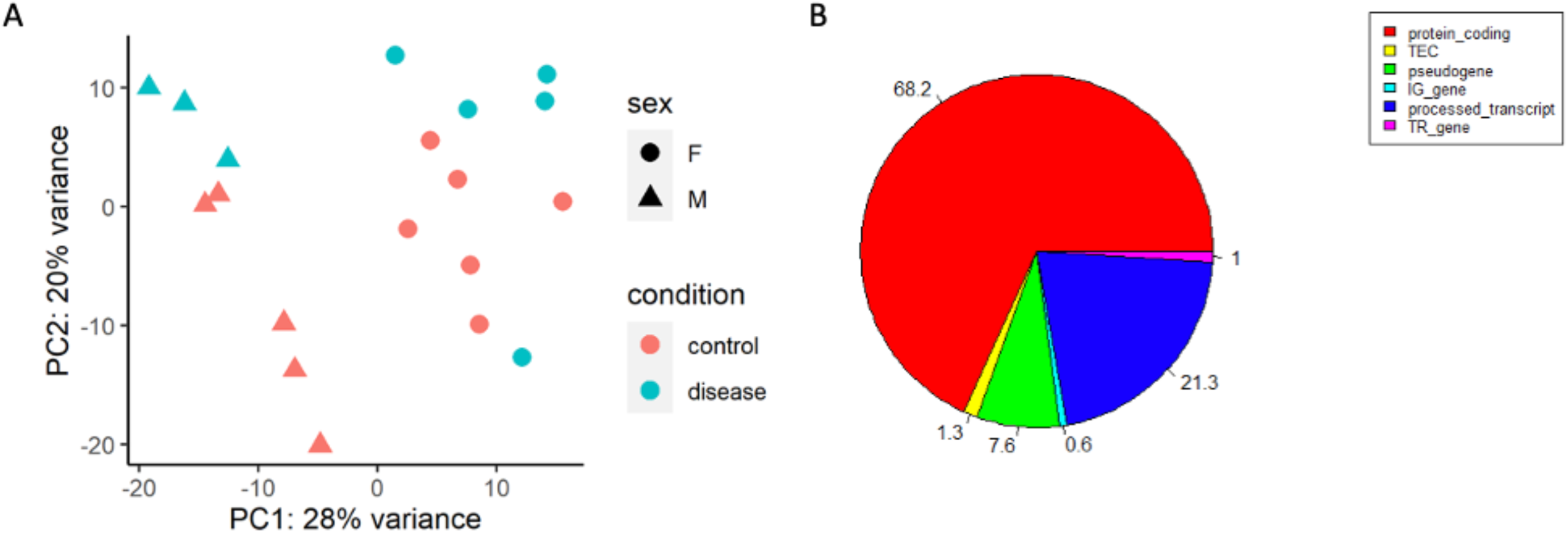
RNAseq Blood Library QC. (**A**) Principal component analysis with adjustment for sex identifies moderate separation of patient and control samples. (**B**) Mapping of reads to transcripts identified a high percentage corresponding to protein-coding genes (68.2%)

**Supplemental Figure 15:**
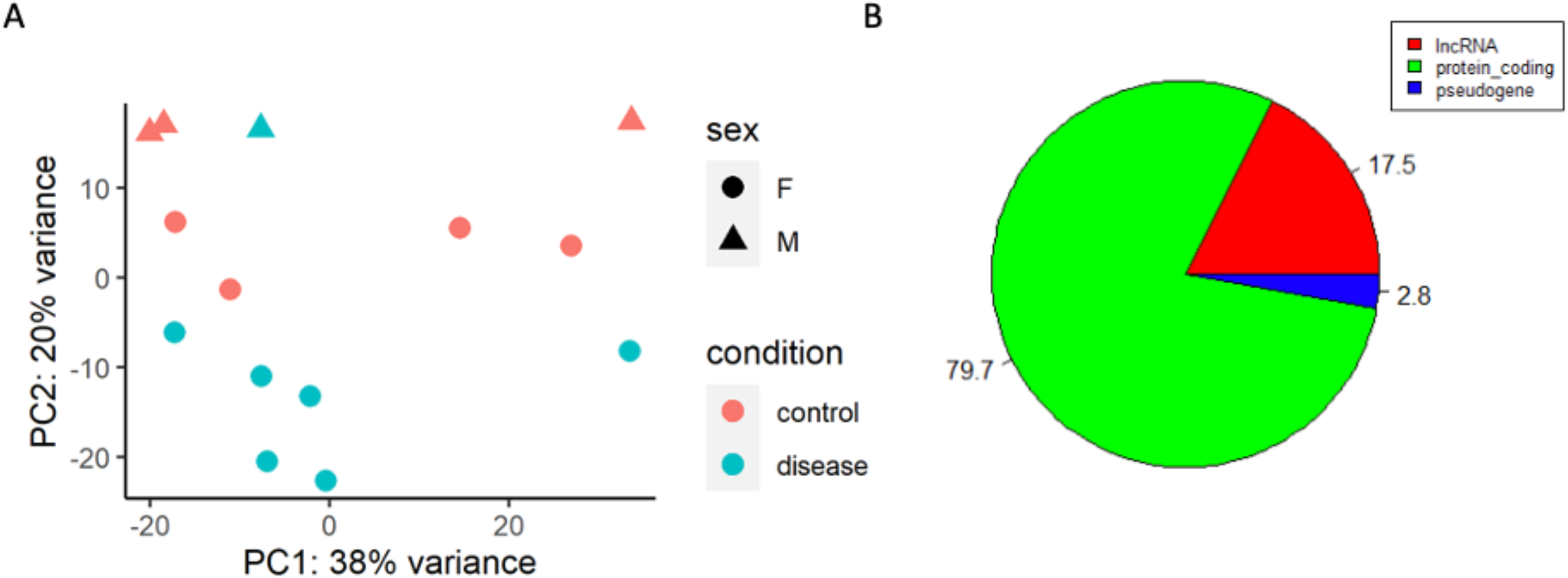
RNAseq Fibroblast Library QC. (**A**) Principal component analysis with adjustment for sex identifies moderate separation of patient and control samples. (**B**) Mapping of reads to transcripts identified a high percentage corresponding to protein-coding genes (79.7%)

**Supplemental Figure 16:**
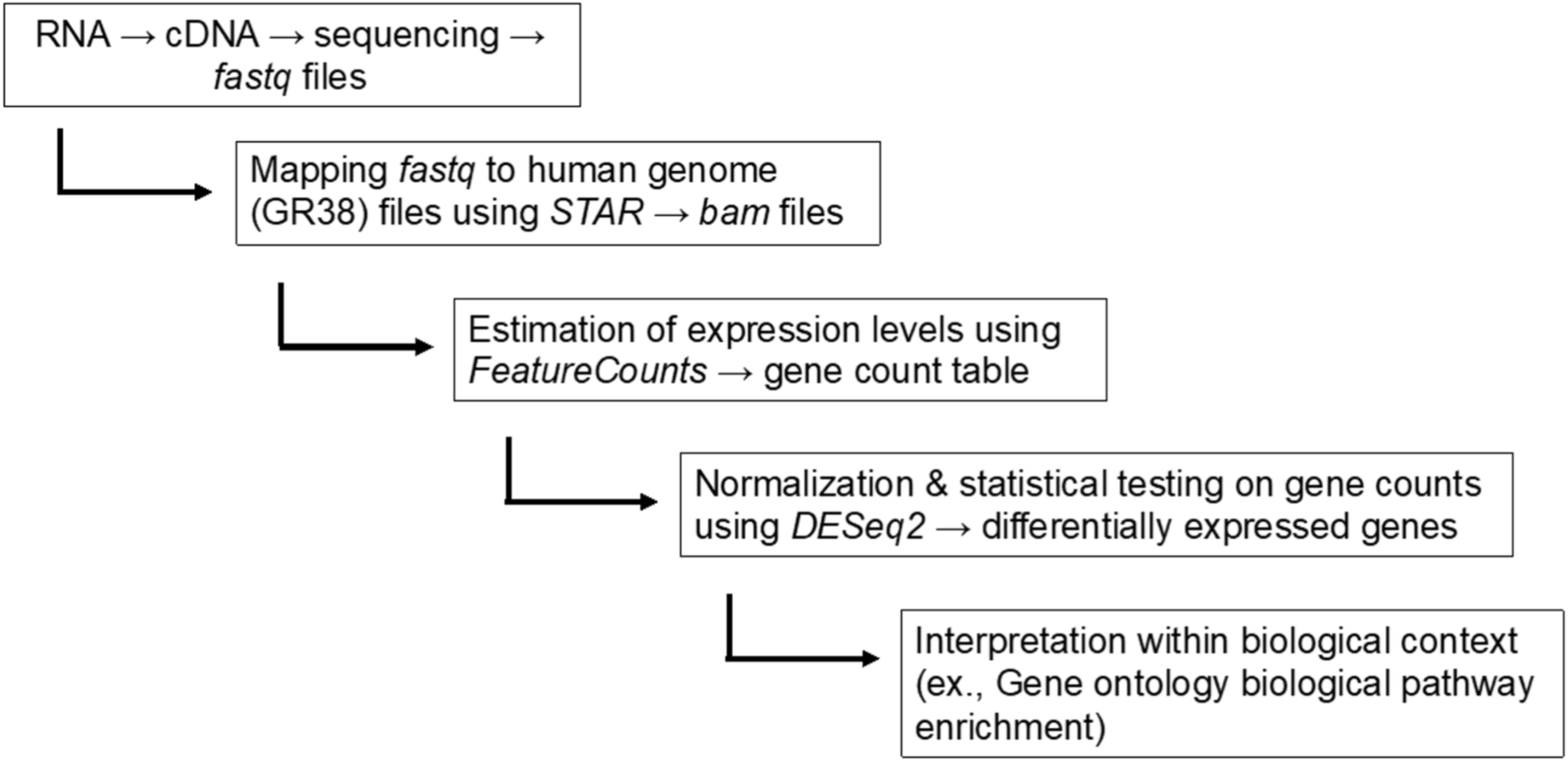
Arboleda Lab best practice RNAseq analysis pipeline.

**Supplemental Figure 17:**
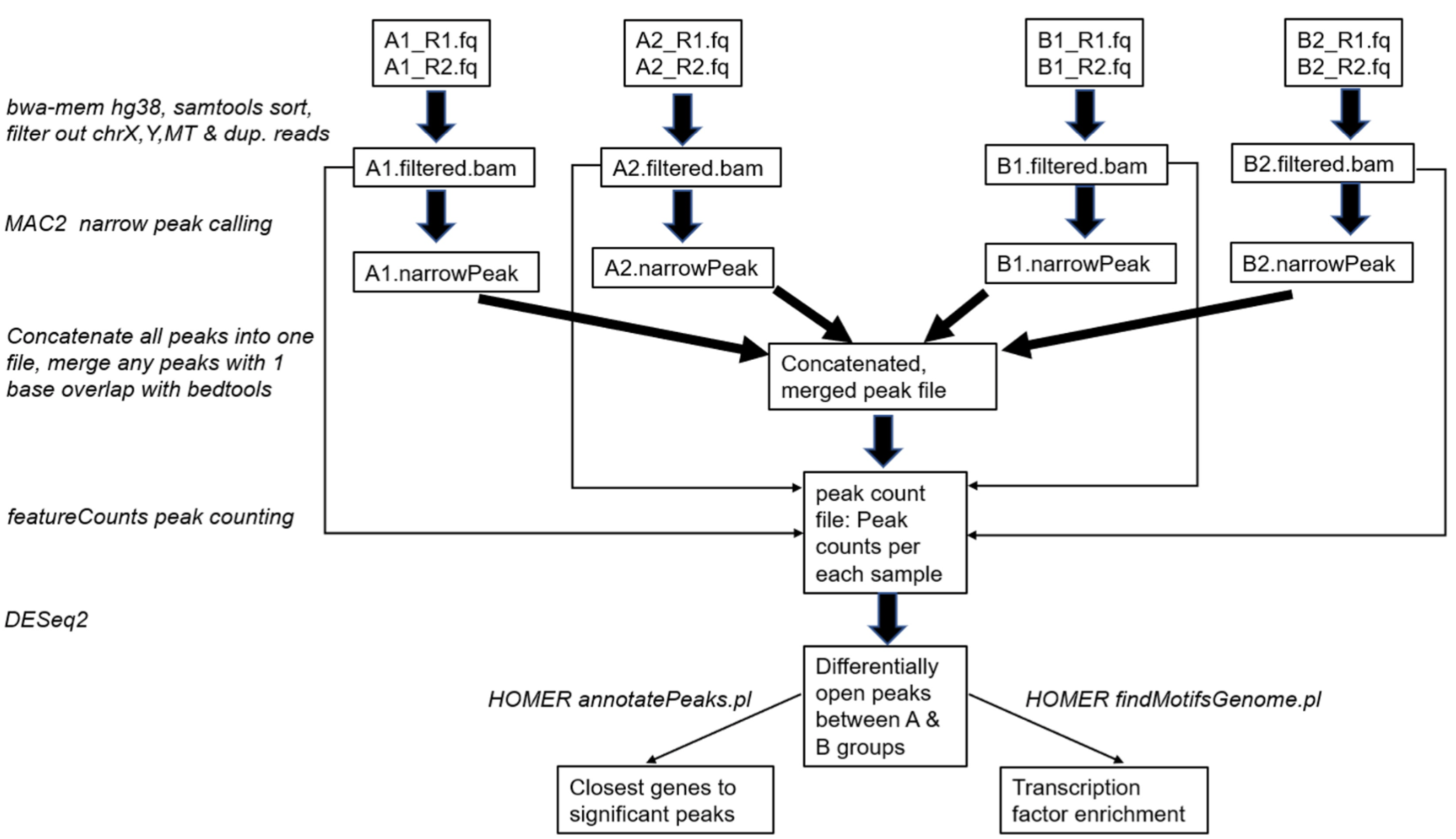
Arboleda Lab best practice ATACseq analysis pipeline.

### SUPPLEMENTAL METHODS

#### Whole Cell Lysis Protein Extraction

Cells were harvested in ice-cold 1X cell lysis buffer (10X, Cell Signaling #9803S) containing 1X Halt™ Protease and Phosphatase Inhibitor Cocktail (100X, Thermo Scientific™ #PI78442). Soluble lysate fractions were isolated by constant agitation at 4℃ for 30 minutes at 1000 rpm and then centrifugation at 20,000 × *g*, for 20 min at 4 °C.

#### Cytoplasmic and Nuclear Extractions

1 million cells were harvested into a 1.5ml eppendorf tube and centrifuged at 500 ×g for 5 min to form a pellet. Cells were washed with ice cold PBS with 0.5mM Sodium Butyrate and centrifuged to form a pellet. Cytoplasmic and nuclear extraction was conducted using the Thermo Scientific CE/NER Kit (#78833) according to protocol. Further details are in Supplemental Methods.

#### Histone Extraction

Histones were extracted as per established protocol (1).

#### Protein Quantification

Protein was quantified using the Pierce BCA Protein Assay Kit (Thermo Scientific #23225), according to protocol.

#### Cytoplasmic and Nuclear Extraction Protocol (Extended)

1 million cells were harvested into a 1.5ml eppendorf tube and centrifuged at 500 ×g for 5 min to form a pellet. Cells were washed with ice cold PBS with 0.5mM Sodium Butyrate and centrifuged at 500 × g for 2 min to form a pellet. After centrifugation, the supernatant was discarded. Ice-cold CERI with 1X Halt™ Protease and Phosphatase Inhibitor Cocktail (100X, Thermo Scientific™ #PI78442) was added to each cell pellet, vortexed on the highest setting for 15 seconds, and incubated on ice for 10 minutes. Then, ice-cold CERII with 1X Halt™ Protease and Phosphatase was added to each tube, vortexed for 5 seconds, and then incubated on ice for 1 minute. The tubes were vortexed for 5 seconds and centrifuged at max speed (16,000 ×g) for 5 min at 4 °C. Immediately after, the supernatant (cytoplasmic extract) was transferred to their respective clean pre-chilled tube and stored at -80°C. The cell pellets were then suspended in ice cold NER with 1X Halt™ Protease and Phosphatase and vortexed for 5 seconds. The samples were then incubated on ice for 40 minutes, with 15 second vortexes at every 10 minute interval. After 40 minutes, the 1.5ml eppendorf tubes were centrifuged at max speed (16,000 ×g) for 10 min at 4 °C. Immediately after, the supernatant (nuclear extract) was transferred to their respective clean pre-chilled tube and stored at -80°C. Protein was quantified using the Pierce BCA Protein Assay Kit (Thermo Scientific #23225).

#### RNAseq Bioinformatic Analysis

An outline of the analysis pipeline is shown in Figure S16. Samples were demultiplexed after sequencing, and fastq sequencing files were processed using established pipelines in the Arboleda lab. Raw read quality, adaptor content, and duplication rates were assessed with *FastQC v0.11.8*.(1) Raw reads were then aligned against the Gencode human genome version hg38 (GR38) version 31 using STAR *2.7.0e* (2) with default parameters. Gene counts from raw reads were generated using featureCounts *1.6.5* (3) from the Subread package. Only reads that uniquely mapped to exons of a gene were counted per gene. Differential expression analysis was completed using DESeq2 *v1.24.0* (*4*), adjusting for sample gender. Genes with a p-adjusted (Benjamini-Hochberg) less than 0.05 were classified as significantly differentially expressed (Wald’s test), and fold changes (4) were shrunk using approximate posterior estimation for GLM coefficients. Gene ontology (5, 6) over-enrichment tests were completed using clusterProfiler *v3.12.0* (*7*) by submitting differentially expressed genes against all genes from the Gencode hg38 annotation, version 31. Gene ontologies were classified as significantly enriched when p-adjusted (Benjamini-Hochberg) was less than 0.05 (hypergeometric test). *HOMER v4.9 findMotifs.pl* (*8*) was used to identify enrichment of motifs within differentially expressed genes’ promoters. Differentially expressed genes were compared against HOMER’s default human, RefSeq-based promoter set, which yielded de novo and known motif enrichments for motif lengths of 8, 10, and 12. Sequencing tracks were visualized using Integrated Genomic Viewer (IGV 2.9.4) and GVIZ.

#### ATACseq Bioinformatic Analysis

An outline of the ATACseq Bioinformatic pipeline is shown in Figure S17. Quality of reads were assessed using FastQC. Raw reads were then aligned to GENCODE Human genome version hg38 (GR38) version 31 using BWA-MEM (9). BAM files were then sorted, indexed, filtered against chrX, chrY, and MT reads using *SAMtools*. *Picard* tools were then used to generate insert size histograms and remove duplicates from BAM files. Narrow peaks from each sample were called using MACS2 callpeak (10); any peak that overlapped by at least one base was then merged using BEDtools merge (11) *merge*. Reads overlapping merged peaks were counted using featureCounts (3). Peaks were also annotated using annotatePeaks.pl from *HOMER* (*8*). *DESeq2* was used to identify differentially open peaks between disease and control samples, adjusting for sample gender. Peaks with p-adjusted value (Benjamini-Hochberg) less than 0.05 were classified as significantly differentially open (Wald’s test), and fold changes were shrunk using approximate posterior estimation for GLM coefficients. Significant peaks were identified as promoter peaks if their distance from their respective closest gene was less than 1kb upstream or 2kb downstream relative to the gene transcription start site. Gene ontology over-enrichment tests were completed using clusterProfiler by submitting the closest genes to significantly differentially open peaks against all genes from the Gencode hg38 annotation, version 31. Gene ontologies were classified as significantly enriched when p-adjusted (Benjamini-Hochberg) was less than 0.05 (hypergeometric test). Enriched de novo and known motifs were identified in significant peaks using findMotifsGenome.pl from HOMER (8). Significant peaks were identified as promoter peaks if their distance from their respective closest gene was less than 1kb upstream or 2kb downstream relative to the gene transcription start site. Sequencing tracks were visualized using Integrated Genomic Viewer (IGV 2.9.4) and GVIZ.

### SUPPLEMENTAL TABLE LEGENDS

**Supplemental Table 1: Bohring-Opitz Syndrome (BOS) Patient Information and Assays**

Bohring-Opitz Syndrome (BOS) patients included in our assays are identified with their patient IDs. We include mutation annotation, age, and sex, and outline the multi-omics assays that each patient’s samples were used in.

**Supplemental Table 2: Control Sample Information and Assays**

Control samples used in our assays are shown with their age and sex. We outline the multi-omics assays that each control’s samples were used in.

**Supplemental Table 3: BOS Patient Summary Table**

We show summary data (age, sex, mutation) for BOS patients in our cohort that are included in each -omics assay.

**Supplemental Table 4: Expression of Key Genes in Tissues and Cell Lines**

Transcripts per million (TPM) gene expression are summarized for key genes of interest. We obtained data for gene expression in relevant tissues from GTEx and ProteinAtlas, and data for cell lines from ProteinAtlas.

**Supplemental Table 5: RT-qPCR Primers**

PrimeTime™ RT-qPCR Primers were purchased from IDT to assay ASXL1 expression. Pre-designed primer pairs and fluorescently labeled 5′ nuclease probe that were used in our assays are listed. These are two primer sets for ASXL1, each targeting a different exonic range. and one set targeting B-actin (ACTB) which was used as a control.

**Supplemental Table 6: List of Antibodies**

Antibodies used in this paper are shown here with their purchasing information.

**Supplemental Table 7: RNAseq Fibroblast QC and Mapping Statistics**

BOS patient and control -derived fibroblasts were used in RNAseq. Here, we show FastQC and mapping statistics for each sample.

**Supplemental Table 8: RNAseq Blood QC and Mapping Statistics**

BOS patient and control blood were used in RNAseq. Here, we show FastQC and mapping statistics for each sample.

**Supplemental Table 9: ASXL1 Pathogenic and Wild-type Allele Counts in BOS patients.**

RNA-seq read counts for the ASXL1 gene in BOS patients was determined. Each read was designated pathogenic if it included the truncating mutation associated with that patient, and wild-type if it did not.

**Supplemental Table 10: Significant Differentially Expressed Genes in BOS Fibroblasts Identified through RNA-seq**

DESeq2 was used to analyze RNAseq gene counts from our cohort of BOS and control fibroblast samples. Samples were adjusted for the co-variate of sex based on principal component analysis. Genes were considered significant if Bonferroni adjusted p < 0.05 and are listed here with their adjusted p value, log_2_Fold Change, and gene annotations.

**Supplemental Table 11: Significant Differentially Expressed Genes in BOS Blood Identified through RNA-seq**

DESeq2 was used to analyze RNAseq gene counts from our cohort of BOS and control blood samples. Samples were adjusted for the co-variate of sex based on principal component analysis. Genes were considered significant if Bonferroni adjusted p < 0.05 and are listed here with their adjusted p value, log_2_Fold Change, and gene annotations.

**Supplemental Table 12: Gene Ontology Analysis for BOS Blood RNA-seq**

Gene ontology (GO) over-enrichment tests were performed using all significant differentially expressed genes in BOS blood samples from our RNA-seq assay. P-values shown are the probability of seeing at least x number of genes out of the total input genes annotated to a particular GO term, given the proportion of genes in the whole genome that are annotated to that GO term. We filtered for gene ontologies with padj < 0.05 for significance.

**Supplemental Table 13: Gene Ontology Analysis for BOS Fibroblast RNA-seq**

Gene ontology (GO) over-enrichment tests were performed using all significant differentially expressed genes in BOS fibroblast samples from our RNA-seq assay. P-values shown are the probability of seeing at least x number of genes out of the total input genes annotated to a particular GO term, given the proportion of genes in the whole genome that are annotated to that GO term. We filtered for gene ontologies with padj < 0.05 for significance.

**Supplemental Table 14: Motif Enrichment Analysis of RNAseq DEGs in BOS Fibroblast**

HOMER (Hypergeometric Optimization of Motif EnRichment) was used to analyze motif enrichment of our BOS RNA-seq fibroblast data using both known motif and de novo methods.

**Supplemental Table 15: ATACseq Fibroblast QC and Mapping Statistics**

BOS patient and control -derived fibroblasts were used in ATAC-seq. Here, we show FastQC, fragment length, and mapping statistics for each sample.

**Supplemental Table 16: Significant Differentially Expressed Genes in BOS Fibroblasts Identified through ATAC-seq**

DESeq2 was used to analyze ATACseq gene counts from our cohort of BOS and control fibroblast samples. Samples were adjusted for the co-variate of sex based on principal component analysis. Genes were considered significant if Bonferroni adjusted p < 0.05 and are listed here with their adjusted p value, log_2_Fold Change, and gene annotations.

**Supplemental Table 17: Gene Ontology Analysis for BOS Fibroblast ATAC-seq**

Gene ontology (GO) over-enrichment tests were performed using all significant differentially expressed genes in BOS fibroblast samples from our ATAC-seq assay. P-values shown are the probability of seeing at least x number of genes out of the total input genes annotated to a particular GO term, given the proportion of genes in the whole genome that are annotated to that GO term. We filtered for gene ontologies with padj < 0.05 for significance.

**Supplemental Table 18: Motif Enrichment Analysis of ATACseq DEGs in BOS Fibroblast**

HOMER (Hypergeometric Optimization of Motif EnRichment) was used to analyze motif enrichment of our BOS ATAC-seq fibroblast data using both known motif and de novo methods.

**Supplemental Table 19: Overlapping DEGs Identified through Chromatin Accessibility and Transcriptional Dysregulation in BOS Fibroblasts**

Bonferroni adjusted p < 0.05 significant differentially expressed genes (DEGs) from our fibroblast RNA-seq dataset (Table S10) and our fibroblast ATAC-seq (Table S16) datasets were integrated to identify common dysregulated genes. We identified 25 DEGs common between the two tissue types, with 21 of these genes dysregulated in the same direction.

**Supplemental Table 20: Significant Differentially Methylated CpG Sites in BOS Blood Identified through DNA methylation**

Significant CpG sites between BOS patient blood and control blood were identified if they were below FDR < 0.05. The absolute difference between the means of the β value for BOS patients versus controls for each CpG was calculated to obtain the delta beta (Δβ) value. We filtered for highly differentially methylated sites (|delta beta (Δβ)| > 10%) using linear regression modeling. Genes were considered significant if below FDR < 0.05 and are listed here with their adjusted p value, Δβ, and gene annotations.

**Supplemental Table 21: Overlapping DEGs between RNAseq and DNA methylation analysis of BOS blood**

Bonferroni adjusted p < 0.05 significant differentially expressed genes (DEGs) from our blood RNA-seq (Table S11) and our blood DNA methylation (Table S19) datasets were integrated to identify common dysregulated genes. We identified 341 DEGs common between the two assay types.

**Supplemental Table 22: Gene Ontology Analysis for BOS Blood RNAseq and DNA Methylation Integration**

Integration of DEGs from our BOS RNAseq blood (Table S11) and BOS blood DNA methylation (Table S19) were analyzed using clusterProfiler *v3.12.0* to identify gene set enrichments. clusterProfiler *v3.12.0* uses all genes from the Gencode hg38 annotation, version 31, as background. We filtered for gene ontologies with padj < 0.05 for significance.

**Supplemental Table 23: Dysregulation of Wnt Signaling Genes in BOS RNAseq and ATACseq Datasets**

Log_2_FoldChange and Bonferroni adjusted p values for RNAseq and ATAC-seq analyses of BOS patient samples are listed for Wnt signaling genes (KEGG pathway).

